# Selection of sequence motifs and generative Hopfield-Potts models for protein families

**DOI:** 10.1101/652784

**Authors:** Kai Shimagaki, Martin Weigt

## Abstract

Statistical models for families of evolutionary related proteins have recently gained interest: in particular pairwise Potts models, as those inferred by the Direct-Coupling Analysis, have been able to extract information about the three-dimensional structure of folded proteins, and about the effect of amino-acid substitutions in proteins. These models are typically requested to reproduce the one- and two-point statistics of the amino-acid usage in a protein family, *i.e*. to capture the so-called residue conservation and covariation statistics of proteins of common evolutionary origin. Pairwise Potts models are the maximum-entropy models achieving this. While being successful, these models depend on huge numbers of *ad hoc* introduced parameters, which have to be estimated from finite amount of data and whose biophysical interpretation remains unclear. Here we propose an approach to parameter reduction, which is based on selecting collective sequence motifs. It naturally leads to the formulation of statistical sequence models in terms of Hopfield-Potts models. These models can be accurately inferred using a mapping to restricted Boltzmann machines and persistent contrastive divergence. We show that, when applied to protein data, even 20-40 patterns are sufficient to obtain statistically close-to-generative models. The Hopfield patterns form interpretable sequence motifs and may be used to clusterize amino-acid sequences into functional sub-families. However, the distributed collective nature of these motifs intrinsically limits the ability of Hopfield-Potts models in predicting contact maps, showing the necessity of developing models going beyond the Hopfield-Potts models discussed here.

## I. INTRODUCTION

Thanks to important technological advances, exemplified in particular by next-generation sequencing, biology is currently undergoing a deep transformation towards a data-rich science. As an example, the number of available protein sequences deposited in the Uniprot database was about 1 million in 2004, crossed 10 millions in 2010, and 100 millions in 2018, despite an important reorganization of the database in 2015 to reduce redundancies and thus limit database size [1]. On the contrary, proteins with detailled experimental knowledge are contained in the manually annotated SwissProt sub-database of Uniprot. While their number remained almost constant and close to 500,000 over the last decade, the knowledge about these selected proteins has been continuously extended and updated.

This fastly growing wealth of data is presenting both a challenge and an opportunity for data-driven modeling approaches. It is a challenge, because for less than 0.5% of all known protein sequences at least some knowledge going beyond sequence is available. Applicability of standard supervised machine-learning approaches is thus frequently limited. However, more importantly, it is an opportunity since protein-sequence databases like Uniprot are not large sets of unrelated random sequences, but contain structured, functional proteins resulting from natural evolution.

In particular, protein sequences can be classified into so-called *homologous protein families* [2]. Each family contains protein sequences, which are believed to share common ancestry in evolution. Such homologous sequences typically show very similar three-dimensional folded structures, and closely related biological functions. Put simply, they can be seen as equivalent proteins in different species, or in different pathways of the same species. Despite this high level of structural and functional conservation, homologous proteins may differ in more than 70-80% of their amino-acids. Detecting homology between a currently uncharacterized protein and a well-studied one [3, 4] is therefore the most important means for computational sequence annotation, including protein-structure prediction by homology modeling [5, 6].

To go beyond such knowledge transfer, we can explore the observable sequence variability between homologous proteins, since it contains on its own important information about the evolutionary constraints acting on proteins to conserve their structure and function [7]. Typically very few random mutations do actually destabilize proteins or interrupt their function. Some positions need to be highly conserved, while others are permissive for multiple mutations. Observing sequence variability across entire homologous protein families, and relating them to protein structure, function, and evolution, is therefore an important task [8].

Over the last years, *inverse statistical physics* [9] has played an increasing role in solving this task. Methods like Direct-Coupling Analysis (DCA) [10, 11] or related approaches [12, 13] allow for predicting protein structure [14, 15], mutational effects [16–18] and protein-protein interactions [19]. However, many of these methods depend on huge numbers of typically *ad hoc* introduced parameters, making these methods data-hungry and susceptible to overfitting effects.

In this paper, we describe an attempt to substantially reduce the amount of parameters, and to select them systematically using sequence data. Despite this parameter reduction, we aim at so-called *generative* statistical models: samples drawn from these models should be statistically similar to the real data, even if similarity is evaluated using statistical measures, which were not used to infer the model from data.

To this aim, we first review in Sec. II some important points about protein-sequence data, maximum-entropy models of these data in general, and profile and DCA models in particular. In Sec. III, we introduce a way for rational selection of so-called sequence motifs, which generalizes maximum-entropy modeling. The resulting Hopfield-Potts models are mapped to Restricted Boltzmann Machines (recently introduced independently for proteins in [20]) in Sec. IV to enable efficient model inference and interpretation of the model parameters. Sec. V is dedicated to the application of this scheme to some exemplary protein families. The Conclusion and Outlook in Sec. VI is followed by some technical appendices.

## II. A SHORT SUMMARY: SEQUENCE FAMILIES, MAXENT MODELS AND DCA

To put our work into the right context, we need to review shortly some published results about the statistical models of protein families. After introducing the data format, we summarize the maximum-entropy approach typically used to justify the use of Boltzmann distributions for protein families, together with some important shortcomings of this approach. Next we give a concise overview over two different types of maximum-entropy models – profile models and direct-coupling analysis – which are currently used for protein sequences. For all cases we discuss the strengths and limitations, which have motivated our current work.

### A. Sequence data

Before discussing modeling strategies, we need to properly define what type of data is used. Sequences of homologous proteins are used in the form of *multiple-sequence alignments* (MSA), *i.e*. rectangular matrices 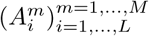. Each of the rows *m* = 1, …, *M* of this matrix contains one aligned protein sequence 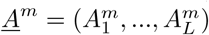 of length *L*. In the context of MSA, *L* is also called the alignment width, *M* its depth. Entries in the matrix come from the alphabet 𝒜= {−, *A, C*, …, *Y*} containing the 20 natural amino acids and the alignment gap “–”. Throughout this paper, the size of the alphabet will be denoted by *q* = 21. In practice we will use a numerical version of the alphabet, denoted by 1, …, *q*, but we have to keep in mind that variables are categorical variables, *i.e*. there is no linear order associated to these numerical values.

The Pfam database [2] currently (release 32.0) lists almost 18,000 protein families. Statistical modeling is most successful for large families, which contain between 10^3^ and 10^6^ sequences. Typical lengths span the range *L* = 30−500.

### B. Maximum-entropy modeling

The aim of statistical modeling is to represent each protein family by a function *P* (*<u>A</u>*), which assigns a probability to each sequence *<u>A</u>* ∈ *𝒜*^*L*^, *i.e*. to each sequence formed by *L* letters from the amino-acid alphabet 𝒜. Obviously the number of sequences even in the largest MSA is much smaller than the number *q*^*L*^ _−_ 1 of a priori independent parameters characterizing *P*. So we have to use clever parameterizations for these models.

A commonly used strategy is based on the maximum-entropy (MaxEnt) approach [21]. It start from any number *p* of observables,

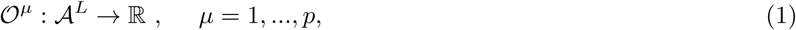

which assign real numbers to each sequence. Only the values of these observables for the sequences in the MSA (*<u>A</u>*^*m*^) go into the MaxEnt models. More precisely, we require the model to reproduce the empirical mean of each observable over the data:

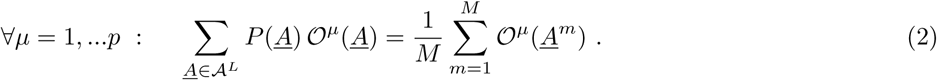

In a more compact notation, we write ⟨ 𝒪^*µ*^⟩_*P*_ = ⟨ 𝒪^*µ*^⟩_MSA_. Besides this consistency with the data, the model should be as unconstrained as possible. Its entropy has therefore to be maximized,

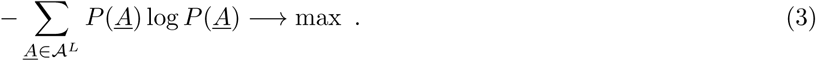

Imposing the constraints in Eq. (2) via Lagrange multipliers *λ*_*µ*_, *µ* = 1, …, *p*, we immediately find that *P* (*<u>A</u>*) assumes a Boltzmann-like exponential form

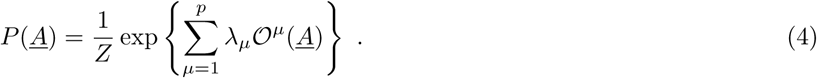

Model inference consists in fitting the Lagrange multipliers such that Eqs. (2) are satisfied. The partition function *Z* guarantees normalization of *P*.

MaxEnt relates observables and the analytical form of the probability distribution, but it does not provide any rule how to select observables. Frequently prior knowledge is used to decide, which observables are “important” and which not. More systematic approaches therefore have to address at least the following two questions:

- Are the selected observables *sufficient*? In the best case, model *P* becomes *generative, i.e*. sequences *<u>A</u>* sampled from *P* are statistically indistinguishable from the natural sequences in the MSA (*<u>A</u>*^*m*^) used for model learning. While this is hard to test in full generality, we can select observables *not* used in the construction of the model, and check if their averages in the model and over the input data coincide.
- Are the selected observables *necessary*? Would it be possible to construct a parameter-reduced, thus more parsimonious model of same quality? This question is very important due to at least two reasons: (a) the most parsimonious model would allow for identifying a minimal set of evolutionary constraints acting on proteins, and thus offer deep insight into protein evolution; and (b) a reduced number of parameters would allow to reduce overfitting effects, which result from the limited availability of data (*M ≪ q*^*L*^).

While there has been promising progress in the first question, cf. the next two subsections, our work attempts to approach both questions simultaneously, thereby going beyond standard MaxEnt modeling.

To facilitate the further discussion, two important technical points have to be mentioned. First, MaxEnt leads to a family of so-called exponential models, where the exponent in Eq. (4) is *linear* in the Lagrange multipliers *λ*_*µ*_, which parameterize the family. Second, MaxEnt is intimately related to maximum likelihood. When we postulate Eq. (4) for the mathematical form of model *P* (*<u>A</u>*), and when we maximize the log-likelihood

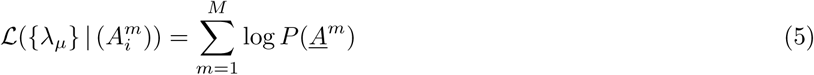

with respect to the parameters *λ*_*µ*_, *µ* = 1, …, *P*, we rediscover Eqs. (2) as the stationarity condition. The particular form of *P* (*<u>A</u>*) guarantees that the likelihood is convex, having only a unique maximum.

### C. Profile models

The most successful approach in statistical modeling of biological sequences are probably *profile models* [22], which consider each MSA column (*i.e*. each position in the sequence) independently. The corresponding observables are simply 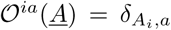, for all positions *i* = 1, …, *L* and all amino-acid letters *a* ∈ 𝒜, with *δ* being the standard Kronecker symbol. These observables thus just ask if in a sequence *<u>A</u>*, amino acid *a* is present in position *i*. Their statistics in the MSA is thus characterized by the fraction

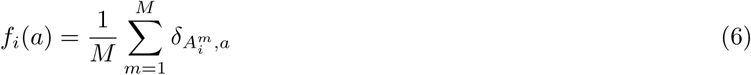

of sequences having amino acid *a* in position *i*. Consistency of model and data requires marginal single-site distributions of *P* to coincide with the *f*_*i*_,

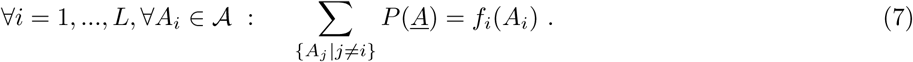

The MaxEnt model results as 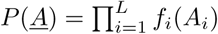, which can be written as a factorized Boltzmann distribution

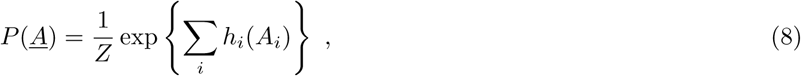

where the local fields equal *h*_*i*_(*a*) = log *f*_*i*_(*a*). Pseudocounts or regularization can be used to avoid infinite negative parameters for amino acids, which are not observed in some MSA column.

Profile models reproduce the so-called *conservation* statistics of an MSA, *i.e*. the heterogenous usage of amino acids in the different positions of the sequence. Conservation of a single or few amino-acids in a column of the MSA is typically an indication of an important functional or structural role of that position. Profile models, frequently in their generalization to profile Hidden Markov Models [3, 4, 23], are used for detecting homology of new sequences to protein families, for aligning multiple sequences, and – using the conserved structural and functional characteristics of protein families – indirectly for the computational annotation of experimentally uncharacterized amino-acid sequences. They are in fact at the methodological basis of the generation of the MSA used here.

Despite their importance in biological sequence analysis, profile models are not generative. Biological sequences show significant correlation in the usage of amino acids in different positions, which are said to *coevolve* [7]. Due to their factorized nature, profile models are not able to reproduce these correlations, and larger sets of observables have to be used to obtain potentially generative sequence models.

### D. Direct-Coupling Analysis

The Direct-Coupling Analysis (DCA) [10, 11] therefore includes also pairwise correlations into the modeling. The statistical model *P* (*<u>A</u>*) is not only required to reproduce the amino-acid usage of single MSA columns, but also the fraction *f*_*ij*_(*a, b*) of sequences having simultaneously amino acid *a* in position *i*, and amino acid *b* in position *j*, for all *a, b ∈ 𝒜* and all 1 *≤ i < j ≤ L*:

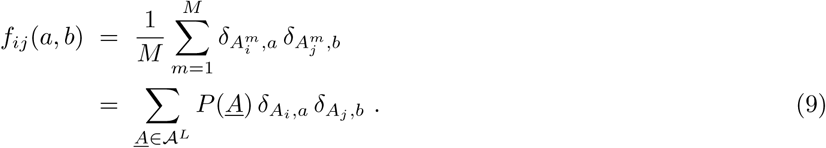

The corresponding observables 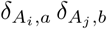 are thus products of pairs of observables used in profile models.

According to the general MaxEnt scheme described before, DCA leads to a generalized *q*-states Potts model

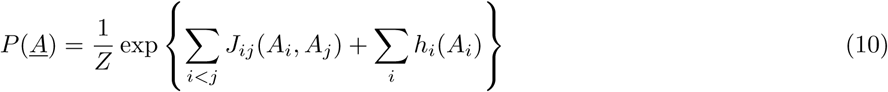

with heterogeneous pairwise couplings *J*_*ij*_(*a, b*) and local fields *h*_*i*_(*a*). The inference of parameters becomes computationally hard, since the computation of the marginal distributions in Eq. (9) requires to sum over *O*(*q*^*L*^) sequences. Many approximation schemes have been proposed, including message-passing [10], mean-field [11], Gaussian [13, 24] and pseudo-likelihood maximization [12, 25] approximations. DCA and related global inference techniques have found widespread applications in the prediction of protein structures, of protein-protein interactions and of mutational effects, demonstrating that amino-acid covariation as captured by the *f*_*ij*_ contains biologically valuable information.

While these approximate inference schemes do not lead to generative models – not even the *f*_*i*_ and the *f*_*ij*_ are accurately reproduced – recently very precise but time-extensive inference schemes based on Boltzmann-machine learning have been proposed [26–29]. Astonishingly, these models do not only reproduce the fitted one- and two-column statistics of the input MSA: also non-fitted characteristics like the three-point statistics *f*_*ijk*_(*a, b, c*) or the clustered organization of sequences in sequence space are reproduced. These observations strongly suggest that pairwise Potts models as inferred via DCA are generative models, *i.e*. that the observables used in DCA – amino-acid occurrence in single positions and in position pairs – are actually defining a (close to) sufficient statistics. In a seminal experimental work [30], the importance of respecting pairwise correlations in amino-acid usage in generating small artificial but folding protein sequences was shown.

However, DCA uses an enormous amount of parameters. There are independent couplings for each pair of positions and amino acids. In case of a protein of limited length *L* = 200, the total number of parameters is close to 10^8^. Very few of these parameters are interpretable in terms of, *e.g*., contacts between positions in the three-dimensional protein fold. We would therefore expect that not all of these observables are really important to statistically model protein sequences. On the contrary, given the limited size (*M* = 10^3^ − 10^4^) of most input MSA, the large number of parameters makes overfitting likely, and quite strong regularization is needed. It would therefore be important to devise parameter-reduced models, as proposed in [31], but without giving up on the generative character of the inferred statistical models.

## III. FROM SEQUENCE MOTIFS TO THE HOPFIELD-POTTS MODEL

Seen the importance of amino-acid conservation in proteins, and of profile models in computational sequence analysis, we keep Eqs. (7), which link the single-site marginals of *P* (*<u>A</u>*) directly to the amino-acid frequencies *f*_*i*_(*a*) in single MSA columns. Further more, we assume that the important observables for our protein-sequence ensemble can be represented as so-called *sequence motifs*

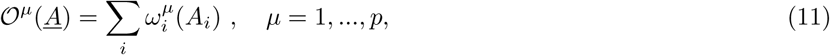

which are linear additive combinations of single-site terms. In sequence bioinformatics, such sequence motifs are widely used, also under alternative names like *position-specific scoring / weight matrices*, cf. [32, 33]. Note that, in difference to the observables introduced before for profile or DCA models, motifs constitute *collective observables* potentially depending on the entire amino-acid sequence.

Let us assume for a moment that these motifs, or more specifically the corresponding *ω*-matrices, are known. We will address their selection later. For any model *P* reproducing the sequence profile, *i.e*. for any model fulfilling Eqs. (7), also the ensemble average of the *𝒪µ* is given,

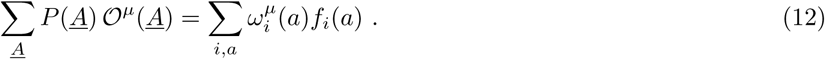

The empirical mean of these observables therefore does not contain any further information about the MSA statistics beyond the profile itself. The key step is to consider also the variance, or the second moment,

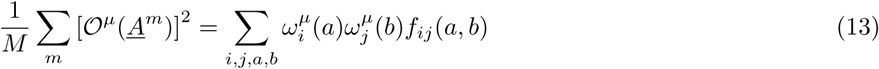

as a distinct feature characterizing the sequence variability in the MSA, which has to be reproduced by the statistical model *P* (*<u>A</u>*). This second moment actually depends on combinations of the *f*_*ij*_, which were introduced in DCA to account for the correlated amino-acid usage in pairs of positions.

The importance of fixing this second moment becomes clear in a very simple example: consider only two positions {1, 2} and two possible letters {*A, B*}, which are allowed in these two positions. Let us assume further that these two letters are equiprobable in these two positions, *i.e. f*_1_(*A*) = *f*_1_(*B*) = *f*_2_(*A*) = *f*_2_(*B*) = 1*/*2. Assume further a single motif to be given by *ω*_1_(*A*) = *ω*_2_(*A*) = 1*/*2, *ω*_1_(*B*) = *ω*_2_(*B*) = − 1*/*2. In this case, the mean of 𝒪 equals zero. We further consider two cases:

- *Uncorrelated positions:* In this case, all words *AA, AB, BA, BB* are equiprobable. The second moment of 𝒪 thus equals 1/2.
- *Correlated positions:* As a strongly correlated example, only the two words *AA* and *BB* are allowed. The second moment of *𝒪* thus equals 1.

We conclude that an increased second moment (or variance) of these additive observables with respect to the uncorrelated case corresponds to the preference of combinations of letters or entire words; this is also the reason why they are the called sequence motifs.

Including therefore these second moments as conditions into the MaxEnt modeling, our statistical model takes the shape

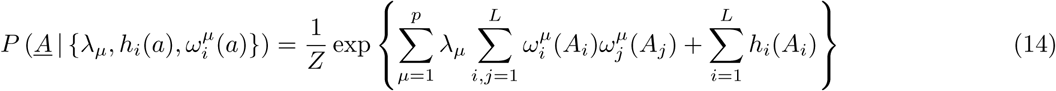

with Lagrange multipliers *λ*_*µ*_, *µ* = 1, …, *p*, imposing means (13) to be reproduced by the model, and *h*_*i*_(*a*), *i* = 1, …, *L, a* ∈ *𝒜*, to impose Eqs. (7).

### A. The Hopfield-Potts model: from MaxEnt to sequence-motif selection

As mentioned before, an important limitation of MaxEnt models is that they assume certain observables to be reproduced, but they do not offer any strategy, how these observables have to be selected. In the case of Eq. (14), this accounts in particular to optimizing the values of the Langrange parameters *λ*_*µ*_ to match the ensemble averages over *P* (*<u>A</u>*) with the sample averages Eq. (13) over the input MSA. As mentioned before, this corresponds also to *maximizing the log-likelihood* of these parameters given MSA and the *ω*-matrices describing the motifs,

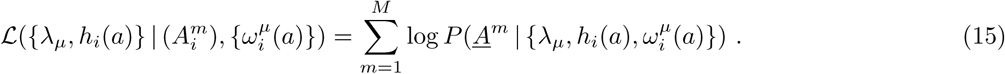

The important, even if quite straight forward *step from MaxEnt modeling to Motif Selection* is to optimize the likelihood also over the choice of all possible *ω*-matrices as parameters. To remove degeneracies, we absorbe the Lagrange multipliers *λ*_*µ*_ into the PSSM *ω*^*µ*^, and introduce

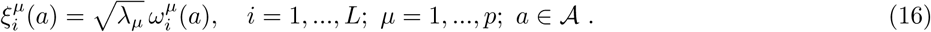

The model in Eq. (14) thus slightly simplifies into

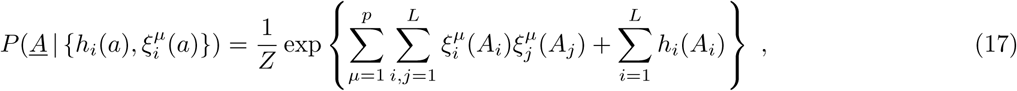

with parameters, which have to be estimated by maximum likelihood:

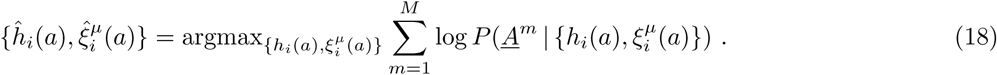

Our model becomes therefore the standard *Hopfield-Potts model*, which has been introduced in [31] in a mean-field treatment, and the sequence motifs equal, up to the rescaling in Eq. (16), the patterns in the Hopfield-Potts model. The mean-field treatment of [31] has both advantages and disadvantages with respect to our present work: on one hand, the largely analytical mean-field solution allows to relate the Hopfield-Potts patterns *ξ*^*µ*^ to the eigenvectors of the Pearson-correlation matrix of the MSA, and their likelihood contributions to a function of the corresponding eigenvalues. This is, in particular, interesting since not only the eigenvectors corresponding to large eigenvalues were found to contribute – as one might expect from the apparent similarity to principal-component analysis (PCA) – but also the smallest eigenvalues lead to large likelihood contributions. However, the mean-field treatment leads to a non-generative model, which does not even reproduce precisely the single-position frequencies *f*_*i*_(*a*). The aim of this paper is to re-establish the generative character of the Hopfield-Potts model by more accurate interefence schemes, without loosing too much of the interpretability of the mean-field approximation.

The model in Eq. (17) contains now an exponent, which is non-linear in the parameters *ξ*^*µ*^. As a consequence, the likelihood is not convex any more, and possibly many local likelihood maxima exist. This is also reflected by the fact that any *p*-dimensional orthogonal transformation of the *ξ*^*µ*^ leaves the probability distribution *P* (*<u>A</u>*) invariant, thus leading to an equivalent model.

## IV. INFERENCE AND INTERPRETATION OF HOPFIELD-POTTS MODELS

### A. The Hopfield-Potts model as a Restricted Boltzmann Machine

The question how many and which patterns are needed for generative modeling therefore cannot be answered properly within the mean-field approach. We therefore propose a more accurate inference scheme based on *Restricted Boltzmann Machine* (RBM) learning [34, 35], exploiting an equivalence between Hopfield models and RBM originally shown in [36]. To this aim, we first perform *p* Hubbard-Stratonovich transformations to linearize the exponential in the *ξ*^*µ*^,

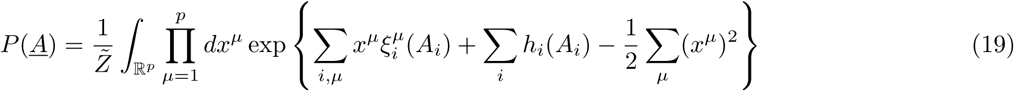

with 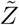 containing the normalizations both of the Gaussian integrals over the new variables *x*^*µ*^, and the partition function of Eq. (14). The distribution *P* (*<u>A</u>*) can thus be understood as a marginal distribution of

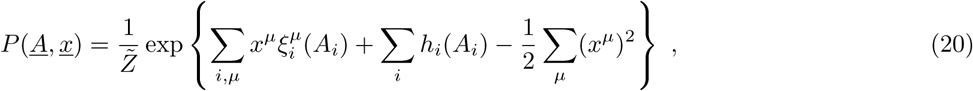

which depends on the so-called *visible variables <u>A</u>* = (*A*_1_, …, *A*_*L*_) and the *hidden (or latent) variables <u>x</u>* = (*x*^1^, …, *x*^*p*^). It takes the form or a particular RBM, with a quadratic confining potential for the *x*^*µ*^: The important point is that couplings in the RBM form a bipartite graph between visible and hidden variables, cf. Fig. 1. RBM may have more general potentials *u*_*µ*_(*x*^*µ*^) confining the values of the new random variables *x*^*µ*^. This fact has been exploited in [20] to cope with the limited number of sequences in the training MSA. However, in our work we stick to quadratic potentials in order to keep the equivalence to Hopfield-Potts models, and thus the interpretability of patterns in terms of pairwise residue-residue couplings via Eq. (17).

**FIG. 1:**
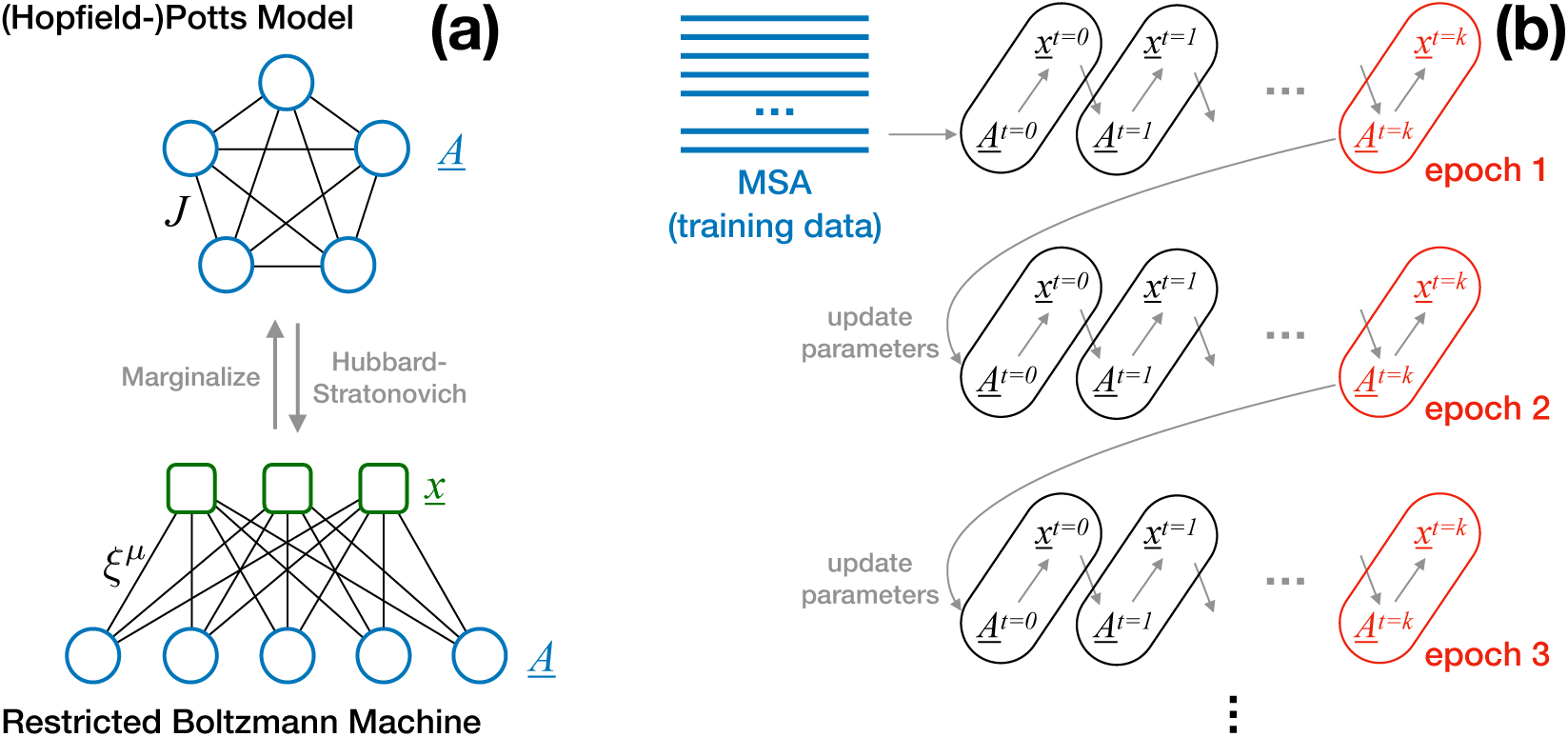
**Panel (a)** represents the (Hopfield-)Potts model as a statistical model for sequences *<u>A</u>* ∈ 𝒜 ^*L*^, typically characterized by a fully connected coupling matrix *J* and local fields *h* (not represented). The model can be transformed into a Restricted Boltzmann Machine (RBM) by introducing Gaussian hidden variables *<u>x</u>* ∈ℝ^*p*^, with *p* being the rank of *J*. Note the bipartite graphical structure of RBM, which causes the conditional probabilities *P* (*<u>A</u>*|*<u>x</u>*) and *P* (*<u>x</u>* |*<u>A</u>*) to factorize. **Panel (b)** shows a schematic representation of Persistent Contrastive Divergence (PCD). Initially the sample is initialized in the training data (the MSA of natural sequences), and then *k* alternating steps of sampling from *P* (*<u>A</u>* |*<u>x</u>*) resp. *P* (*<u>x</u>*| *<u>A</u>*) are performed. Parameters are updated after these *k* sampling steps, and sampling is continued using the updated parameters.

### B. Parameter learning by persistent contrastive divergence

Maximizing the likelihood with respect to the parameters leads, for our RBM model, to the stationarity equations

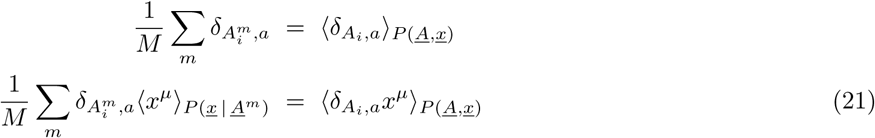

for all *i, a* and *µ*; the difference of both sides equals the gradient of the likelihood in direction of the corresponding parameter. While the first line matches the standard MaxEnt form – sample and ensemble average of an observable have to coincide, the second line contains a mixed sample-ensemble average on its left-hand side. Since the variables *x*^*µ*^ are latent and thus not contained in the MSA, an average over their probability *P* (*<u>x</u>*|*<u>A</u>*^*m*^) conditioned to the sequences *<u>A</u>*^*m*^ in the MSA has to be taken. Having a *P* -dependence on both sides of Eqs. (21) is yet another expression of the non-convexity of the likelihood function.

Model parameters *h*_*i*_(*a*) and 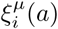 have to be fitted to satisfy the stationarity conditions Eq. (21). This can be done iteratively: starting from arbitrarily initialized model parameters, we determine the difference between the left- and right-hand sides of this equation, and use this difference to update parameters (*i.e*. we perform gradient ascent of the likelihood); each of these update steps is called an *epoch* of learning. A major problem is that the exact calculation of averages over the (*L* + *p*)-dimensional probability distribution *P* is computationally infeasible. It is possible to estimate these averages by Markov Chain Monte Carlo (MCMC) sampling, but efficient implementations are needed since accurate parameter learning requires in practice thousands of epochs. To this aim, we exploit the bipartite structure of RBM: both conditional probabilities *P* (*<u>A</u>*|*<u>x</u>*) and *P* (*<u>x</u>*|*<u>A</u>*) are factorized. This allows us to initialise MCMC runs in natural sequences from the MSA and to sample the *<u>x</u>* and the *<u>A</u>* in alternating fashion. As a second simplification we use *Persistent Contrastive Divergence* (PCD) [37]. Only in the first epoch the visible variables are initialised in the MSA sequences, and each epoch performs only a finite number of sampling steps (*k* for PCD-*k*), cf. Fig. 1.B. Trajectories are continued in a new epoch after parameter updates. If the resulting parameter changes become small enough, PCD will thereby generate close-to-equilibrium sequences, which form an (almost) *i.i.d*. sample of *P* (*<u>A</u>, <u>x</u>*) uncorrelated from the training set used for initialization.

Details of the algorithm, and comparison to the simpler contrastive divergence are given in the Appendix. Further technical details, like regularization, are also delegated to the Appendix.

### C. Determining the likelihood contribution of single Hopfield-Potts patterns

It is obvious, that the total likelihood grows monotonously when increasing the number *p* of patterns *ξ*^*µ*^. It is therefore important to develop criteria, which tell us if patterns are more or less important for modeling the protein family. To this aim we estimate the contribution of single patterns to the likelihood, by comparing the full model with a model, where a single pattern *ξ*^*µ*^ has been removed, while the other *p* −1 patterns and the local fields have been retained. The corresponding normalized change in log-likelihood reads

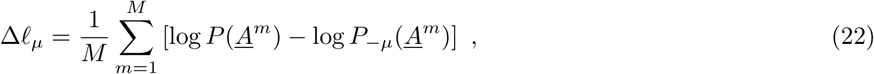

where *P*_*−µ*_ has the same form as given in Eq. (14) for *P*, but with pattern 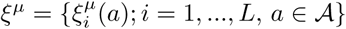 removed. Plugging Eq. (14) into Eq. (22), we find

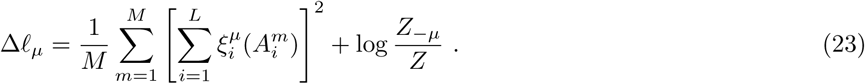

The likelihood difference depends thus on the ratio of the two partition functions *Z* and *Z*_−*µ*_. While each of them is individually intractable due to the exponential sum over *q*^*L*^ sequences, the ratio can be estimated efficiently using importance sampling. We write:

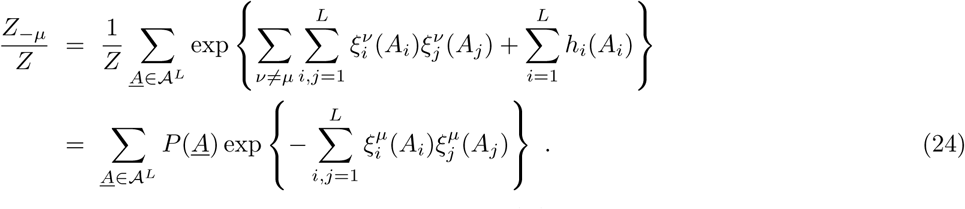

The last expression contains the average of an exponential quantity over *P* (*<u>A</u>*), so estimating the average by MCMC sampling of *P* might appear a risky idea. However, since *P* and *P*_*-µ*_ differ only in one of the *p* patterns, the distributions are expected to overlap strongly, and sufficiently large samples drawn from *P* (*<u>A</u>*) can be used to estimate *Z*_−*µ*_*/Z*. Note that sampling is done from *P*, so the likelihood-contributions of all patterns can be estimated in parallel using a single large sample of the full model.

Once these likelihood contributions are estimated, we can sort them, and identify and interpret the patterns of largest importance in our Hopfield-Potts model.

## V. HOPFIELD-POTTS MODELS OF PROTEIN FAMILIES

To understand the performance of Hopfield-Potts models in the case of protein families, we have analysed three protein families extracted from the Pfam database [2]: the Kunitz/Bovine pancreatic trypsin inhibitor domain (PF00014), the Response regulator receiver domain (PF00072) and the RNA recognition motif (PF00076). They have been selected since they have been used in DCA studies before, in our case RBM results will be compared to the ones of bmDCA, *i.e*. the generative version of DCA based on Boltzmann Machine Learning [29]. MSA are downloaded from the Pfam database [2], and sequences with more than 5 consecutive gaps are removed; cf. App. B for a discussion of the convergence problems of PCD-based inference in case of extended gap stretches. The resulting MSA dimensions for the three families are, in the order given before, *L* = 52*/*112*/*70 and *M* = 10657*/*15000*/*10000. As can be noted, the last two MSA have been subsampled randomly since they were very large, and the running time of the PCD algorithm is linear in the sample size. The MSA for PF00072 was chosen to be slightly larger because of the longer sequences in this family.

In the following sections, results are described in detail for the PF00072 response regulator family. The results for the other protein families are coherent with the discussion; they are moved to the App. B for the seek of conciseness of our presentation.

### A. Generative properties of Hopfield-Potts models

PCD is able, for all values of the pattern number *p*, to reach parameter values satisfying the stationarity conditions Eqs. (21). This is not only true when these are evaluated using the PCD sample propagated via learning from epoch to epoch, but also when the inferred model is resampled using MCMC, i.e. when the right-hand side of Eqs. (21) is evaluated using an *i.i.d*. sample of the RBM.

In the leftmost column of Fig. 2 (Panels (a.1-g.1)), this is shown for the single-site frequencies, *i.e*. for the first of Eqs. (21). The horizontal axis shows the statistics extracted from the original data collected in the MSA, while the vertical axis measures the same quantity in an *i.i.d*. sample extracted from the inferred model *P* (*<u>A</u>, <u>x</u>*). The fitting quality is comparable to the one obtained by bmDCA, as can be seen by comparison with the last panel in the first column of Fig. 2.

**FIG. 2:**
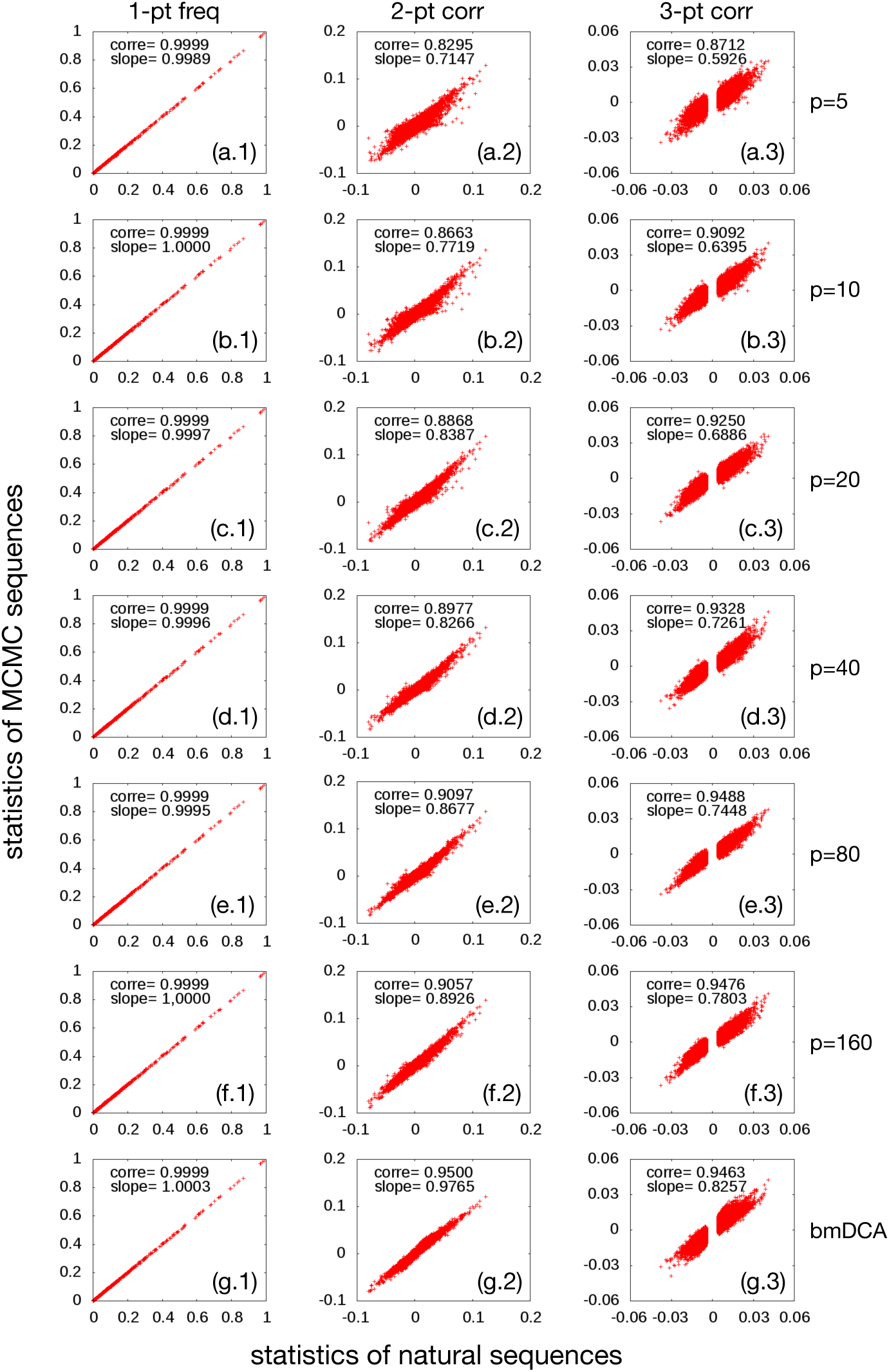
Statistics of natural sequences (PF00072, horizontal axes) vs. MCMC samples (vertical axes) of Hopfield-Potts models for values of *p*∈ {5, 10, 20, 40, 80, 160} and for a full-rank Potts model inferred using bmDCA. The first column (Panels (a.1-g.1)) shows the 1-point frequencies *f*_*i*_(*a*) for all pairs (*i, a*) of sites and amino-acids, the other two column show the connected 2- and 3-point functions *c*_*ij*_(*a, b*) (Panels (a.2-g.2)) and *c*_*ijk*_(*a, b, c*) (Panels (a.3-g.3)). Due to the huge number of combinations for the three-point correlations, only the 100,000 largest values (evaluated in the training MSA) are shown. The Pearson correlations and the slope of the best linear fit are inserted in each of the panels.

The other two columns of the figure concern the *generative* properties of RBM: connected two-point correlations (Panels (a.2-g.2)) and three-point correlations (Panels (a.3-g.3)),

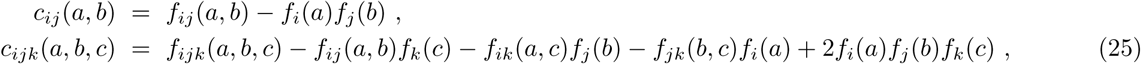

with the three-point frequencies *f*_*ijk*_(*a, b, c*) defined in analogy to Eqs. (6,9). Note that in difference to DCA, already the two-point correlations are not fitted directly by the RBM, but only the second moments related to the Hopfield-Potts patterns. This becomes immediately obvious for the case *p* = 0, where RBM reduce to simple profile models of statistically independent sites, but remains true for all values of *p <* (*q* − 1)*L*. Note also that connected correlations are used, since the frequencies *f*_*ij*_ and *f*_*ijk*_ contain information about the fitted *f*_*i*_, and therefore show stronger agreement between data and model.

The performance of RBM is found to be, up to statistical fluctuations, monotonous in the pattern number *p*. As in the mean-field approximation [31], no evident overfitting effects are observed. Even if not fitted explicitly, as few as *p* = 20 − 40 patterns are sufficient to faithfully reproduce even the non-fitted two- and three-point correlations. This is very astonishing, since only about 1.7-3.5% of the parameters of the full DCA model are used: the *p* patterns are given by *p*(*q −* 1)*L* parameters, while DCA has 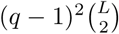 independently inferred couplings. The times needed for accurate inference decrease accordingly: in some cases, a slight decrease in accuracy of bmDCA is observed as compared to RBM with the largest *p*; this could be overcome by iterating the inference procedure for further epochs.

### B. Strong couplings and contact prediction

One of the main applications of DCA is the prediction of contacts between residues in the three-dimensional protein fold, based only on thI:e statistics of homologous sequences. To this aim, we follow [25] and translate *q × q* coupling matrices 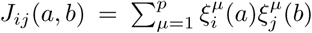 for individual site pairs (*i, j*) into scalar numbers by first calculating their Frobenius norm,

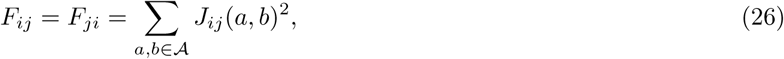

followed by the empirical average-product correction (APC)

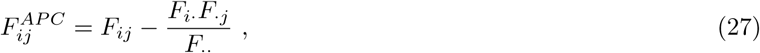

where the · denotes an average over the corresponding index:

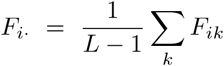

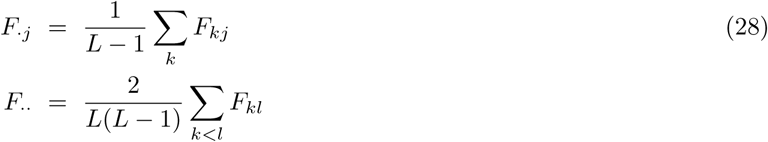

The APC is intended to remove systematic non-functional bias due to conservation and phylogeny. These quantities are sorted, and the largest ones are expected to be contacts.

The results for several values of *p* and for bmDCA are depicted in Fig. 3.a: the positive predictive value (PPV) is the fraction of true positives (TP) among the first *n* predictions, as a function of *n*. TP are defined as native contacts in a reference protein structure (PDB ID 3ilh [38] for PF00072), with a distance cutoff of 8Å between the closest pair of heavy atoms forming each residue. Pairs in vicinity along the peptide chain are not considered in this prediction, since they are trivially in contact: in coherence with the literature standard, Fig. 3 only considers predictions with.*|i - j|* ≥5

**FIG. 3:**
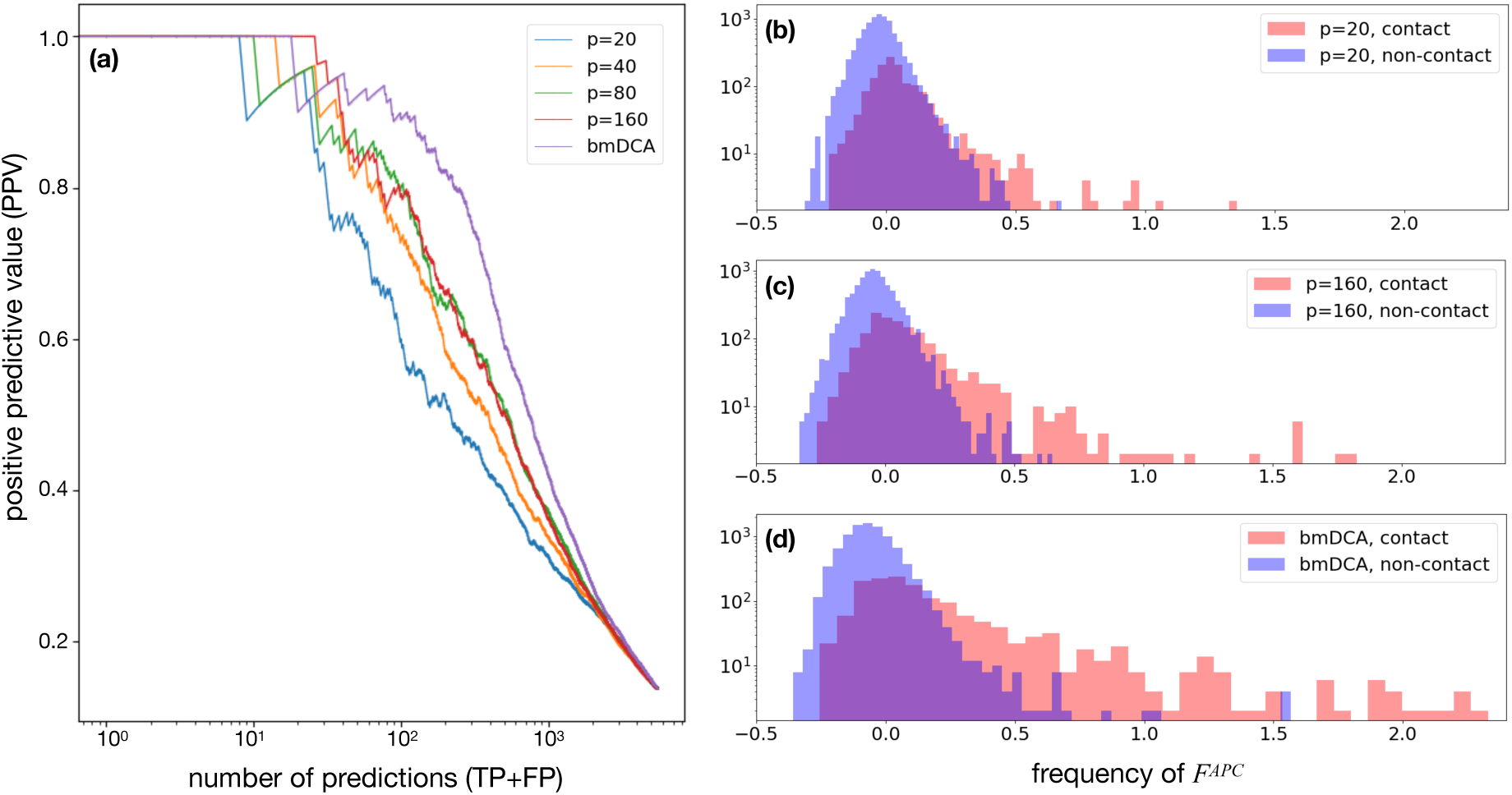
Panel (a) shows the positive predictive value (PPV) for contact prediction as a function of the number of predictions, for various values of the pattern number *p* and for bmDCA. Panels (b-d) show, for *p* = 20, 160 and bmDCA, the distribution of coupling scores 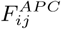. All residue pairs are grouped into contacts (red) and non-contacts (blue). The best contact predictions correspond to the positive tail of the red histogram, which becomes more pronounced when increasing *p* or even going to bmDCA.

Despite the fact, that even for as few as *p* = 20 − 40 patterns the model appears to be generative, *i.e*. non-fitted statistical observables are reproduced with good accuracy, the PPV curves depend strongly on the pattern number *p*. Up to statistically probably insignificant exceptions, we observe a monotonous dependence on *p*, and none of the RBM-related curves reaches the performance of the full-rank *J*_*ij*_ matrices of bmDCA. Even large values of *p*, where RBM have more than 30% of the parameters of the full Potts model, show a drop in performance in contact prediction.

Can we understand this apparent contradiction: similarly accurate reproduction of the statistics, but reduced performance in contact prediction? To this end, we consider in Figs. 3.b-d the histograms of coupling strengths 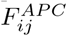 divided into two subpopulations: values for sites *i, j* in contact are represented by red, for distant sites by blue histograms. It becomes evident that the rather compact histogram of non-contacts remains almost invariable with *p* (even if individual coupling values do change), but the histogram of contacts changes systematically: the tail of large 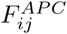 going beyond the upper edge of the blue histogram is less pronounced for small *p*. However, in the procedure described before, these 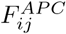-values provide the first contact predictions.

The reduced capacity to detect contacts for small *p* is related to the properties of the Hopfield-Potts model in itself. While the esidue-residue contacts form a sparse graph, the Hopfield-Potts model is explicitly constructed to have a low-rank coupling matrix (*J*_*ij*_(*a, b*)). It is, however, hard to represent a generic sparse matrix by a limited number of possibly distributed patterns. Hopfield-Potts models are more likely to detect distributed sequence signals than localized sparse ones. However, for larger pattern numbers *p* we are able to detect more and more localized signals, thereby improving the contact prediction, until BM and Hopfield-Potts models become equivalent for *p* = (*q* − 1)*L*.

This observation establishes an important limitation to the generative character of Hopfield-Potts models with limited pattern numbers: The applicability of DCA for residue-residue contact prediction has demonstrated that physical contacts in the three-dimensional structure of proteins introduce important constraints on sequence evolution. A perfectly generative model should respect these constraints, and thus lead to a contact prediction being at least as good as the one obtained by full DCA, cf. also the discussion in the outlook of this article.

### C. Likelihood contribution and interpretation of selected sequence motifs

So what do the patterns represent? In Sec. IV C, we have discussed how to estimate the likelihood contribution of patterns, thereby being able to select the most important patterns in our model. Fig. 4 displays the ordered contributions for different values of *p*. We observe that, for small *p*, the distribution becomes more peaked, with few patterns having very large likelihood contributions. For larger *p*, the contributions are more distributed over many patterns, which collectively represent the statistical features of the data set.

**FIG. 4:**
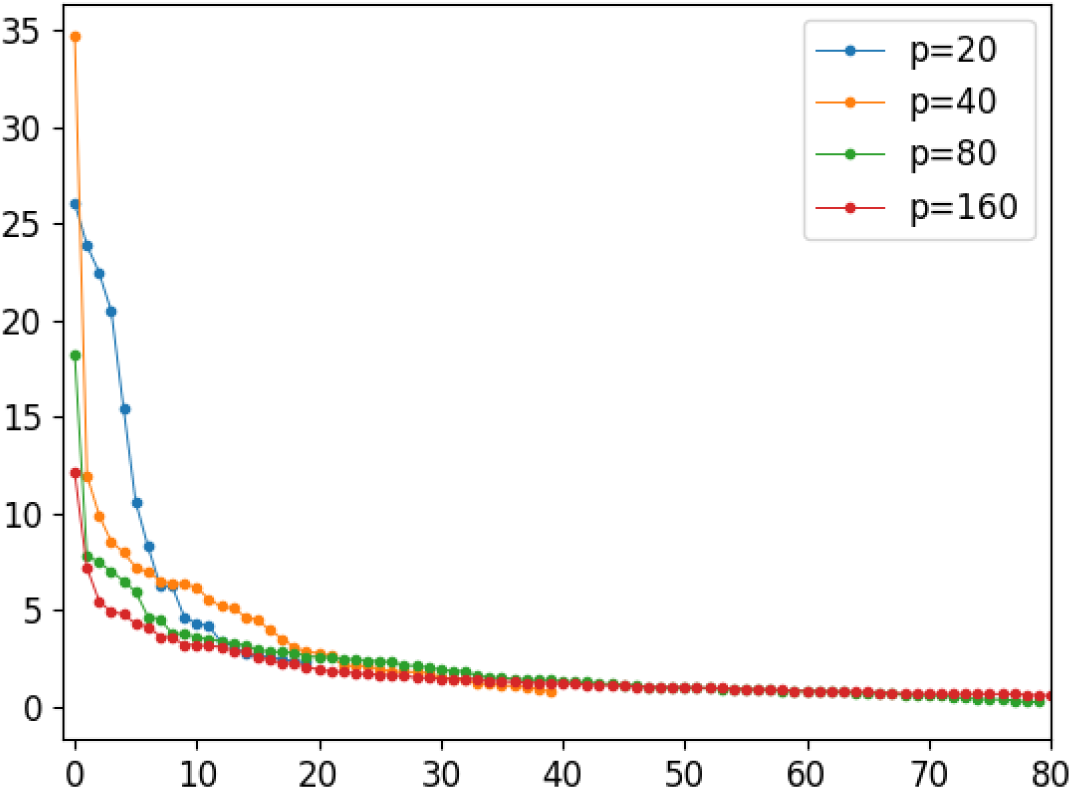
Likelihood contribution of the individual patterns, for pattern numbers *p* = 20, 40, 80, 160.

Fig. 5 represents the first five patterns for *p* = 20. Panels (a.1-e.1) represent the pattern 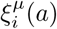 as a sequence logo, a standard representation in sequence bioinformatics. Each site *i* corresponds to one position, the possible amino-acids are shown by their one-letter code, the size of the letter being proportional to 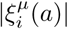, according to the sign of 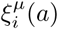, letters are represented above or below the zero line. The alignment gap is represented as a minus sign in an oval shape, which allows to represent its size in the current pattern.

**FIG. 5:**
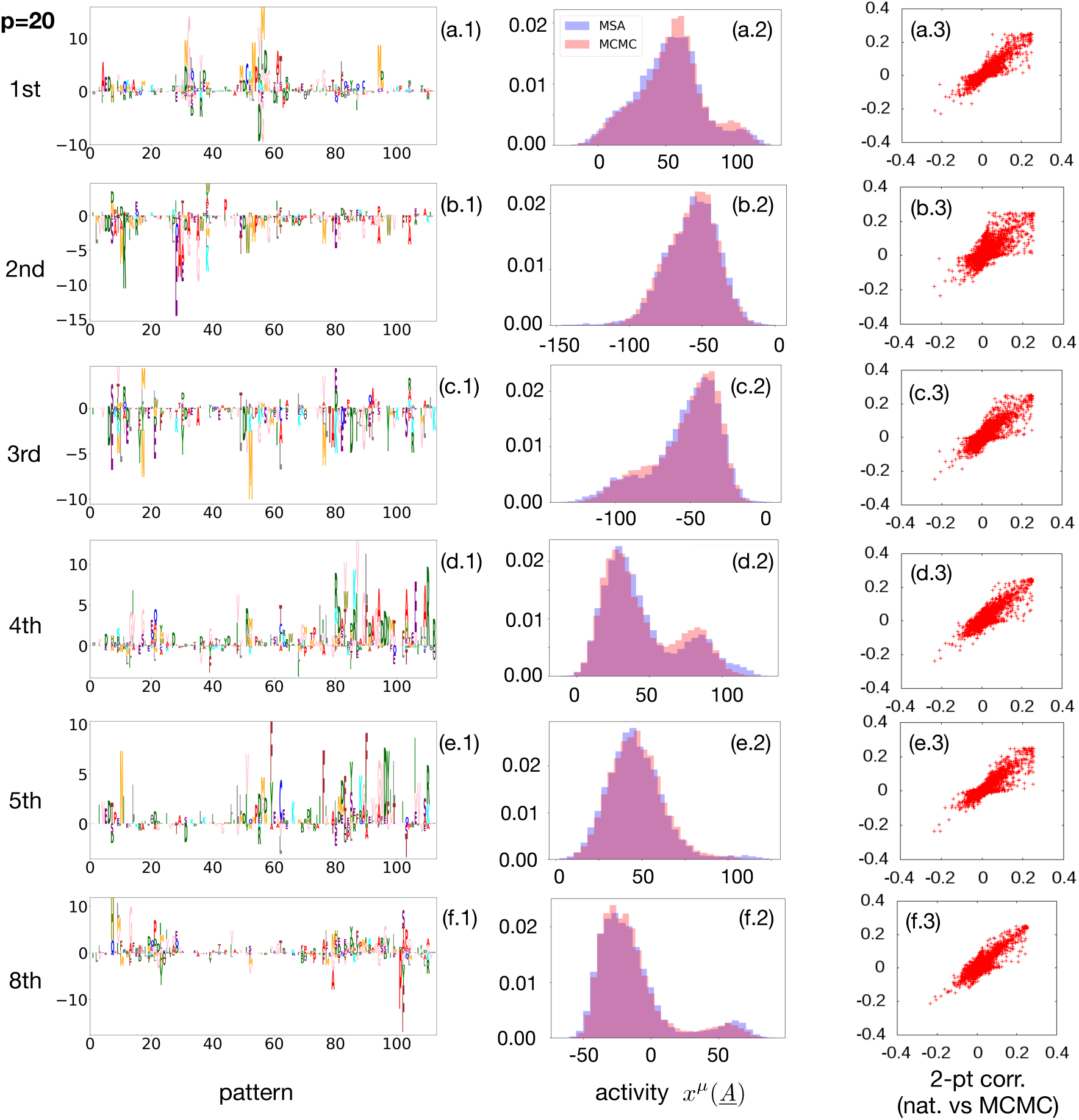
The five patterns of highest likelihood-contribution for *p* = 20, the 8th ranking is added since used later in the text. The left Panels (a.1-f.1) show the patterns in logo representation, the letter-size is given by the corresponding element *ξ*_*i*_(*a*). The middle panels (a.1-f.2) show the distribution of the activities, *i.e*. the projections of sequences onto the patterns. The blue histogram contains the natural sequences from the training MSA, the red histogram sequences sampled by MCMC from the Hopfield-Potts model. The right-hand side (Panels (a.3-f.3)) shows the connected 2-point correlations of the natural data (horizontal axis) vs. data sampled from *P*_*− µ*_(*<u>A</u>*), i.e. a Hopfield-Potts model with one pattern removed. Strong deviations from the diagonal are evident.

Patterns are very distributed, both in terms of the sites and amino-acids with relatively large entries *ξ*_*i*_(*a*). This makes a direct interpretation of patterns without prior knowledge rather complicated. The distributed nature of patterns explains also why they are not optimal in defining localised contact predictions. Rather than identifying contacting residue pairs, the patterns define larger groups of sites, which are connected via a dense network of comparable couplings. However, as we will see in the next section, the sites of large entries in a pattern define functional regions of proteins, which are important in sub-ensembles of proteins of strong (positive or negative) activity values along the pattern under consideration. In particular, we will show that the largest entries may have an interpetation connecting structure and function to sequence in protein sub-families.

The middle column (Panels (a.2-e.2)) shows a histogram of pattern-specific activities of single sequences, *i.e*. of

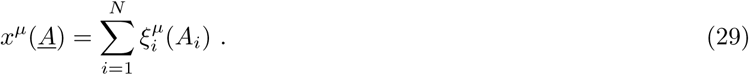

Note that, up to the rescaling in Eq. (16), these numbers coincide with the sequence motifs, introduced in Eq. (11) at the beginning of this article. They also equal the average value of the latent variable *x*^*µ*^ given sequence *<u>A</u>*. The blue histograms result from the natural sequences collected in the training MSA. They coincide well with the red histograms, which are calculated from an *i.i.d*. MCMC sample of our Hopfield-Potts model, including the bimodal structure of several histograms. This is quite remarkable: the Hopfield-Potts model was derived, in the beginning of this work, as the maximum-entropy model reproducing the first two moments of the activities *{x*^*µ*^(*<u>A</u>*^*m*^)*}*_*m*=1…*M*_. Finding higher-order features like bimodality is again an expression of the generative power of Hopfield-Potts models.

Figs. 5.a.3-5.e.3 prove the importance of individual patterns for the inferred model. The panels show the two-point correlations *c*_*ij*_(*a, b*) of the natural data (horizontal axis) vs. the one of samples drawn from the distributions *P*_*−µ*_(*<u>A</u>*), introduced in Eq. (22) as Hopfield-Potts models of *p* − 1 patterns, with pattern *ξ*^*µ*^ removed (vertical axis). The coherence of the correlations is strongly reduced when compared to the full model, which was shown in Fig. 2: removal even of a single pattern has a strong global impact on the model statistics.

### D. Sequence clustering

As already mentioned, some patterns show a clear bimodal activity distribution, *i.e*. they identify two statistically distinct subgroups of sequences. The number of subgroups can be augmented by using more than one pattern, *i.e*. combinations of patterns can be used to cluster sequences.

To this aim, we have selected three patterns (number 6, 13 and 14) with a pronounced bimodal structure from the model with *p* = 20 patterns. In terms of likelihood contribution, they have ranks 8, 4 and 1 in the contributions to the log-likelihood, cf. Fig. 5.

The clustered organization of response-regulator sequences becomes even more evident in the two-dimensional plots characterizing simultaneously two activity distributions. The results for all pairs of the three patterns are displayed in Fig. 6, Panels (a.1-a.3). As a first observation, we see that the main modes of the activity patterns give rise to one dominant cluster. Smaller cluster deviate from the dominant one in a single pattern, but show compatible activities in the other patterns – the two-dimensional plots therefore show typically an L-shaped sequence distribution, and three clusters, instead of the theoretically possible four combinations of activity models. It appears that single patterns identify the particularities of single subdominant sequence clusters.

**FIG. 6:**
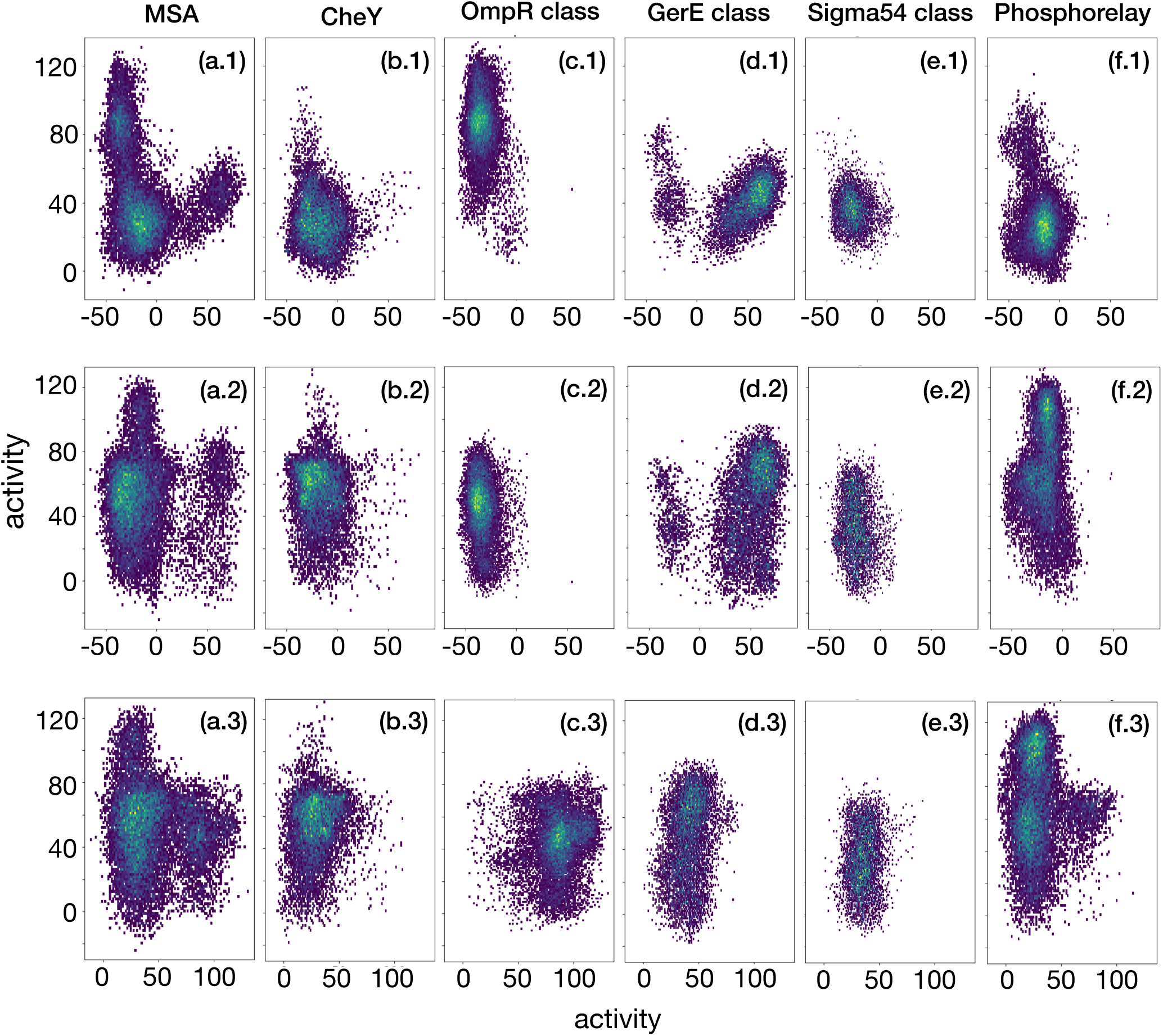
Patterns with multimodal activity distributions for the set of all MSA sequences can be used to cluster sequences. The rows show combinations of patterns 6-13 (Panels (a.1-f.1)), 6-14 (Panels (a.2-f.2)) and 13-14 (Panels (a.3-f.3)). Each sequence corresponds to a density-colored dot. A strongly clustered structure is clearly visible. When dividing the full MSA into functional subclasses, we can relate clusters to subclasses, and thus patterns to biological function.

We have chosen the response-regulator protein-domain family in this paper also due to the fact, that it constitutes a functionally well studied and diversified family. Response regulators are predominantly used in bacterial signaling systems:

- In *chemotaxis*, they appear as single-domain proteins named CheY, which transmit the signal from kinase proteins (activated by signal reception) to flagellar motor proteins, which trigger the movement of the bacteria. CheY proteins can be identified in our MSA as those coming from single-domain proteins, *i.e*. with lengths compatible to the PF00072-MSA width *L* = 112. We have selected a sub-MSA consisting of all proteins with total sequence lengths between 110 and 140 amino acids.
- In *two-component signal transduction* (TCS), response regulators are typically transcription factors, which are activated by signal-receiving histidine sensor kinases. The corresponding proteins contain two or three domains, in particular a DNA-binding domain, which is actually responsible for the transcription-factor activity of the activated response-regulator protein. According to the present DNA-binding domain, these TCS proteins can be subdivided into different classes, the dominant ones are the OmpR, the GerE and the Sigma54-HTH8 classes, we identified three sub-MSA corresponding to these classes by co-occurrence of the DNA-binding domains with the response-regulator domain in the same protein. The different DNA-binding domains are indicative for distinct homo-dimer structures assumed by the active transcription factors; DCA run on the sub-MS identifies their specific sub-family interfaces [39].
- *Phosphorelays* are similar to TCS, but consist of more complex multi-component signaling pathways. In these systems, found in bacteria and plants, response-regulator domains are typically fused to the histidine-sensor kinases. They do not act as transcription factors, but transduct a signal to a phosphotransferase, which finally activates a down-stream transcription factor of the same architecture mentioned in the last paragraph. We identified a class of response regulator domains, which are fused to a histidine kinase domain. In terms of domain architecture and protein length, this subfamily is extremely heterogenous.

Panel columns (a-f) in Fig. 6 show the activities of these five sub-families. It is evident, that distinct sub-MSA fall actually into distinct clusters according to these three patterns:

- The CheY-like single domain proteins (Panels (b.1-b.3)) fall, according to all three patterns, into the dominant mode.
- The OmpR-class transcription factors (Panels (c.1-c.3)) show a distinct distribution of higher activities for the second of the patterns (which actually has the most pronounced bimodal structure, probably due to the fact that the OmpR-class forms the largest sub-MSA). As can be seen in Fig. 6, this pattern has the largest positive entries in the region of positions 80-90 and 100-110. Interestingly, these regions define the interface of OmpR-class transcription-factor homodimerization, cf. [39]. In accordance with this structural interpretation, we also find a periodic structure of period 3-4 of the large entries in the pattern, which reflects the fact that the interface is formed by two helices, which lead to a periodic exposure of amino-acids in the protein surface.
- The GerE-class (Panels (d.1-d.3)) differs in activities in direction of the first pattern, only GerE-class proteins have positive, all other have negative activities. Dominant positive entries are found in regions 5-15 and 100-105, again identifying the homo-dimerization interface, cf. [39].
- The Sigma54 class (Panels (e.1-e.3)) does not show a distinct distribution of activities according to the three selected patterns. It is located together with the CheY-type sequences. However, when examining all patterns, we find that pattern number 5 (ranked 6th according to the likelihood contribution) is almost perfectly discriminating the two.
- Last but not least, the response-regulators fused to histidine kinases in phosphorelay systems (Panels (f.1-f.3)) show a distinct activity distribution according to the third pattern, mixing a part of activities compatible with the main cluster, and others being substantially larger (this mixing results presumably from the previously mentioned heterogenous structure of this sub MSA). Structurally known complexes between response-regulators and histidine phosphotransferases (PDB ID 4euk [40], 1bdj [41]) show the interface located in residues 5-15, 30-32 and 50-55, regions being important in the corresponding pattern. It appears that the pattern selects the particular amino-acid composition of this interface, which is specific to the phosphorelay sub-MSA.

These observations do not only show that the patterns allow for clustering sequences into subMSA, but the discriminating positions in the patterns have a clear biological interpretation. This is very interesting, since the analysis in [39] required a prior clustering of the initial MSA into sub-MSA, and the application of DCA to the individual sub-MSA. Here we have inferred only one Hopfield-Potts model describing the full MSA, and the patterns automatically identify biologically reasonable sub-families together with the sequence patterns characterizing them. The prior knowledge needed in [39] is not needed here; we use it only for the posterior interpretation of the patterns.

It is also important to remember that sequence clustering can also be obtained by a technically simpler PCA (principal-component analysis). PCA is based on the leading eigenvectors of the data-covariance matrix, *i.e*. exclusively on the largest eigenvalues. The potential differences were already discussed in [31] in the context of the mean-field approximation of Hopfield-Potts models. It was shown that not only the eigenvectors with large eigenvalues lead to important contributions in likelihood, but also those corresponding to the smallest eigenvalues. Both tails of the spectrum are thus important for the statistical description of protein-sequence ensembles. A second drawback of PCA as compared to our approach is the non-generative character of PCA. No explicite statistical model is learned, but the data covariance matrix is simply approximated by a low-rank matrix.

## VI. CONCLUSION AND OUTLOOK

In this paper, we have rederived Hopfield-Potts models as statistical models for protein sequences by selection of additive sequence motifs. Statistical sequence models are required to reproduce the first and second moments of the empirical motif distributions (*i.e*. over the MSA of natural sequences). Within a maximum-entropy approach, these motifs are found to be (up to a scaling factor) the Hopfield-Potts patterns defining a network of residue-residue couplings. In addition to the maximum-entropy framework, which is built upon known observables, the Hopfield-Potts model adds a step of variable selection: the probability of the sequence data is maximised over all possible selections of sequence motifs.

The quadratic coupling terms can be linearised using a Hubbard-Stratonovich transformation. When the Gaussian variables introduced in this transformation are interpreted as latent random variables, the Hopfield-Potts model takes the form of a Restricted Boltzmann Machine. This interpretation, originally introduced in [36], allows for the application of efficient inference techniques, like persistent contrastive divergence, and therefore for the accurate inference of the Hopfield-Potts patterns for any given MSA of a homologous protein family.

We find that Hopfield-Potts models acquire interesting generative properties even for a relatively small number of parameters (p=20-40). They are able to reproduce non-fitted properties like higher-order covariation of residues. Also the bimodality observed in the empirical activity distributions (i.e. the projection of the natural sequences onto individual Hopfield-Potts patterns) is not automatically guaranteed when using only the first two moments for model learning, but it is recovered with high accuracy in the activity distributions of artificially sequences sampled from the model. This observation is not only interesting in the context of generative-model learning, but forms the basis of sequence clustering according to interpretable sequence motifs in the main text.

The Hopfield-Potts patterns, or sequence motifs, are typically found to be distributed over many residues, thereby representing global features of sequences. This observation explains, why Hopfield-Potts models tend to loose accuracy in residue-residue contact prediction, as compared to the full-rank Potts models normally used in Direct Coupling Analysis: the sparsity of the residue-residue contact network cannot be represented easily via few distributed sequence motifs, which describe more global patterns of sequence variability. Despite the fact, that Hopfield-Potts models reproduce also non-fitted statistical observables, the loss of accuracy in contact prediction demonstrates that these models are not fully generative, and alternative concepts for parameter reduction should be explored.

Individual sequences from the input MSA can be projected onto the Hopfield-Potts patterns, resulting in sequence-specific activity values. Some patterns show a mono-modal histogram for the protein family. They introduce a dense network of relatively small couplings between positions with sufficiently large entries in the pattern, without dividing the family into subfamilies. These patterns have great similarity to the concept of protein sectors, which was introduced in [42, 43] to detect distributed modes of sequence coevolution. However, the conservation-based reweighting used in determining sectors is not present in the Hopfield-Potts model, and the precise relationship between both ideas remains to be elaborated. Other Hopfield-Potts patterns show a bimodal activity distributions, leading to the detection of functional sub-families. Since these are defined by, e.g., the positive vs. the negative entries of the pattern, the entries of large absolute value in the patterns identify residues, which play a role similar to so-called specificity determining residues [44, 45], *i.e*. residues, which are conserved inside specific sub-families, but vary between sub-families. Both concepts – sectors and specificity-determining residues – emerge naturally in the context of Hopfield-Potts families, even if their precise mathematical definitions differ, and thus also their precise biological interpretations.

These observations open up new ways of parameter reduction in statistical models of protein sequences: the sparsity of contacts, which are expected to be responsible for a large part of localized residue covariation in protein evolution, has to be combined with the low-rank structure of Hopfield-Potts models, which detect distributed functional sequence motifs. However, distributed patterns may also be related to phylogenetic correlations, which are present in the data, cf. [46]. As has been shown recently in a heuristic way [47], the decomposition of sequence-data covariance matrices or couplings matrices into a sum of a sparse and a low-rank matrix can substantially improve contact prediction, if only the sparse matrix is used.

Combining this idea with the idea of generative modeling seems a promising road towards parsimonious sequence models, which in turn would improve parameter interpretability and reduce overfitting effects, both limiting factors of current versions of DCA. In this context, it will also be interesting to explore more general regularization strategies which favor more localized sequence motifs, or Hopfield-Potts patterns, thereby unifying sparse and low-rank inference in a single framework of parameter-reduced statistical models for biological sequence ensembles.

## Acknowledgements

We are particularly grateful to Pierre Barrat-Charlaix, Giancarlo Croce, Carlo Lucibello, Anna-Paola Muntoni, Andrea Pagnani, Edoardo Sarti, Jerome Tubiana and Francesco Zamponi for numerous discussions and assistance with data.

We acknowledge funding by the EU H2020 research and innovation programme MSCA-RISE-2016 under grant agreement No. 734439 InferNet. KS acknowledges an Erasmus Mundus TEAM (TEAM – Technologies for information and communication, Europe - East Asia Mobilities) scholarship of the European Commission in 2017/18, and a doctoral scholarship of the Honjo International Scholarship Foundation since 2018.

## Appendix A: Results for other protein families

The first appendix is dedicated to other protein families. As discussed in the main text, we have analyzed three distinct families, and discussed only one in full detail in the main text. Here we present the major results – generative properties, contact prediction and selected collective variables (patterns) – for two more families. These results show the general applicability of our approach beyond the specific response-regulator family used in the main text.

### 1. Kunitz/Bovine pancreatic trypsin inhibitor domain PF00014

Figs. 7, 8 et 9 display the major results for the PF00014 protein family. PPV curves are calculated using PDB ID 5pti [48].

**FIG. 7:**
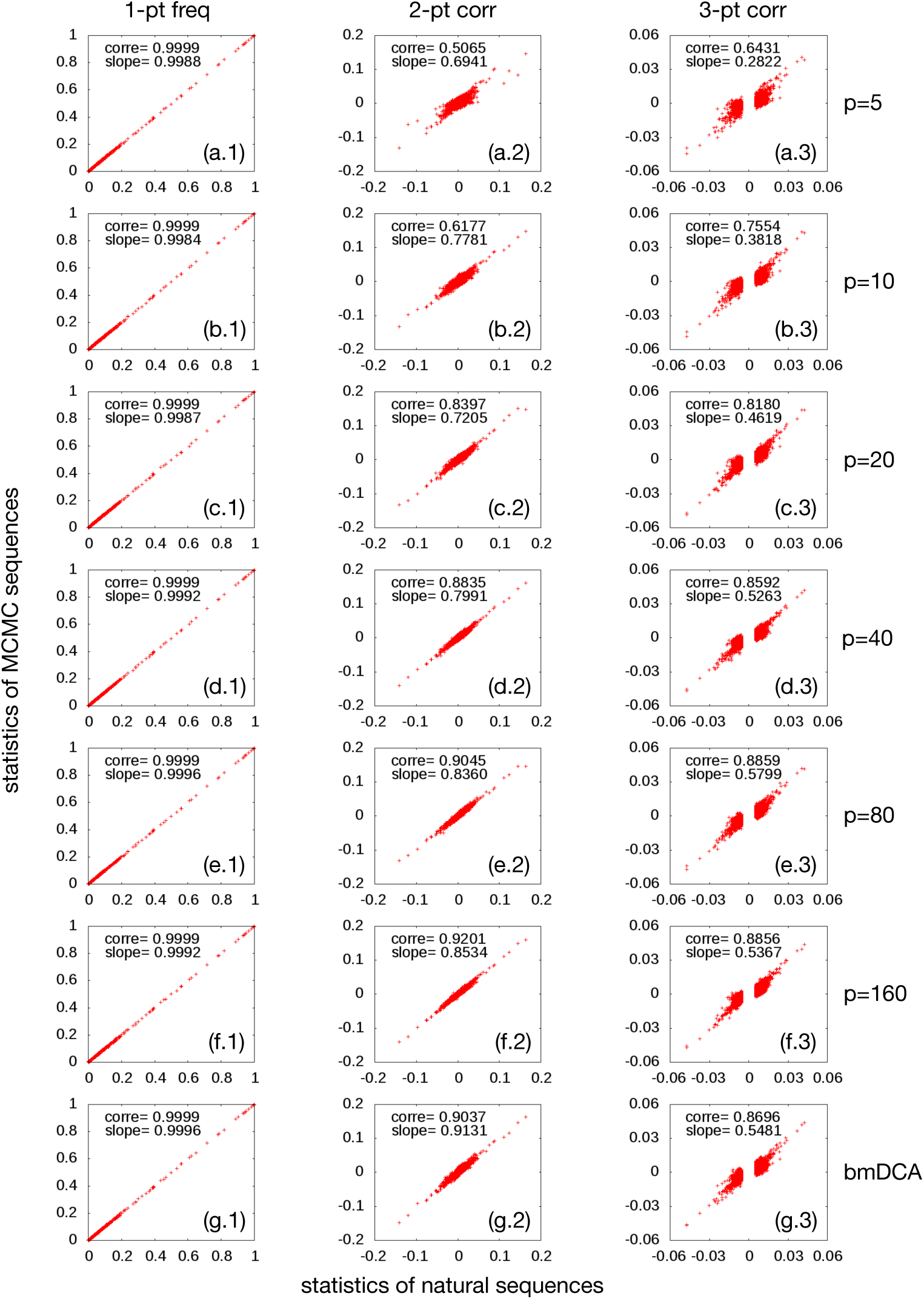
Same as Fig. 2, but for the protein family PF00014.

**FIG. 8:**
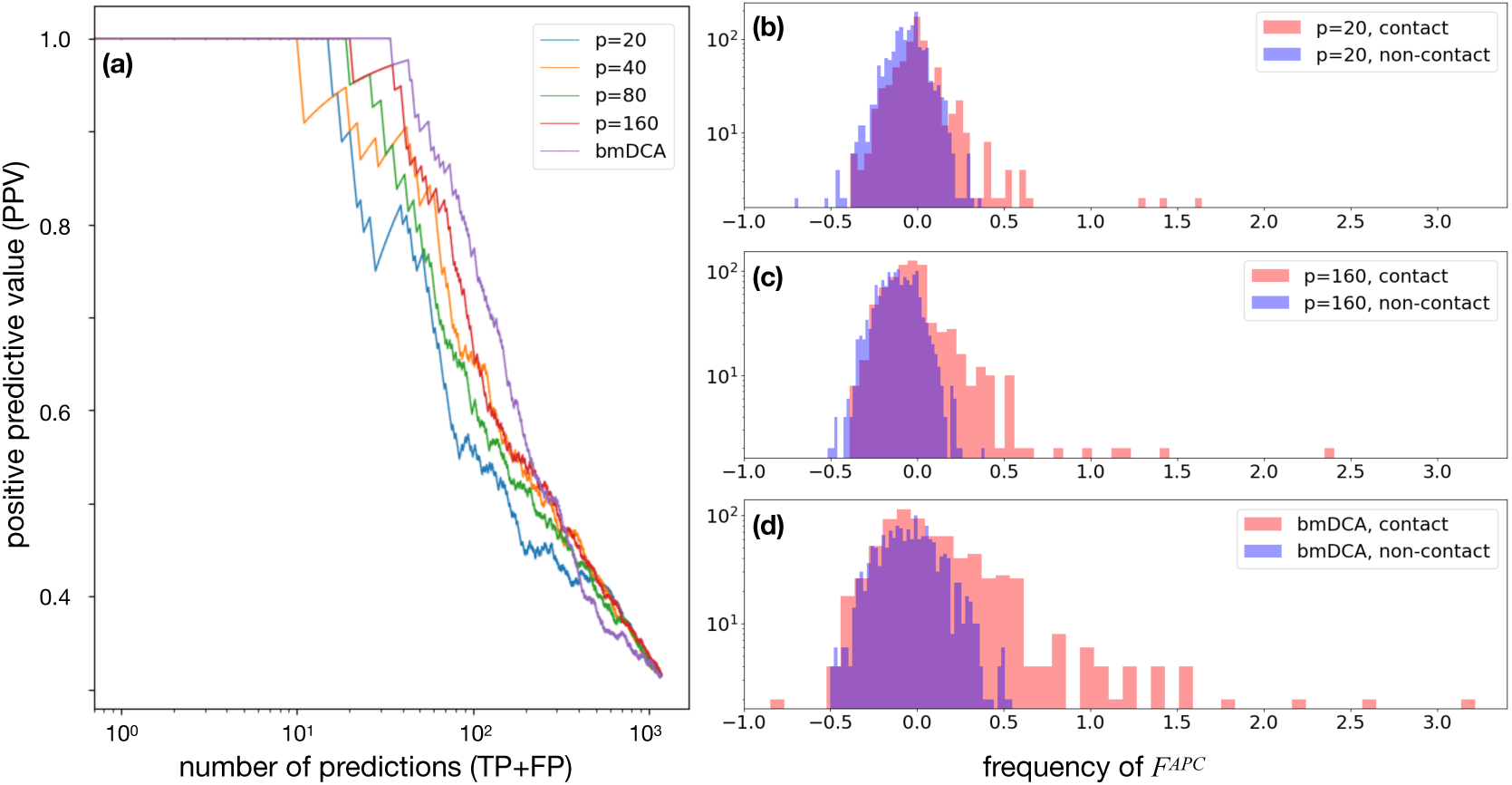
Same as Fig. 3, but for the protein family PF00014.

### 2. RNA recognition motif PF00076

Figs. 10, 11 et 12x display the major results for the PF00076 protein family. PPV curves are calculated using PDB ID 2×1a [49].

**FIG. 9:**
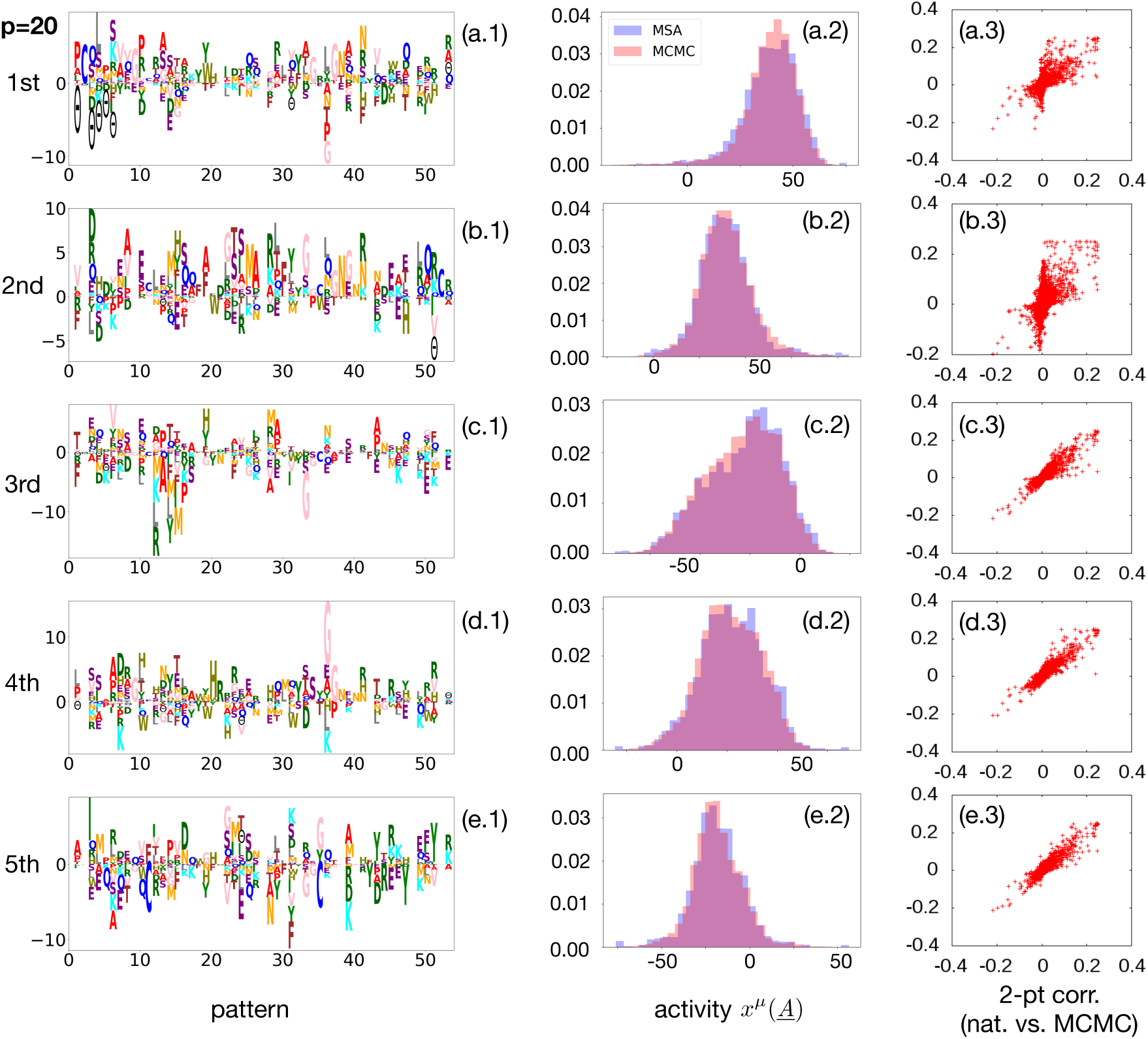
Same as Fig. 5, but for the protein family PF00014.

**FIG. 10:**
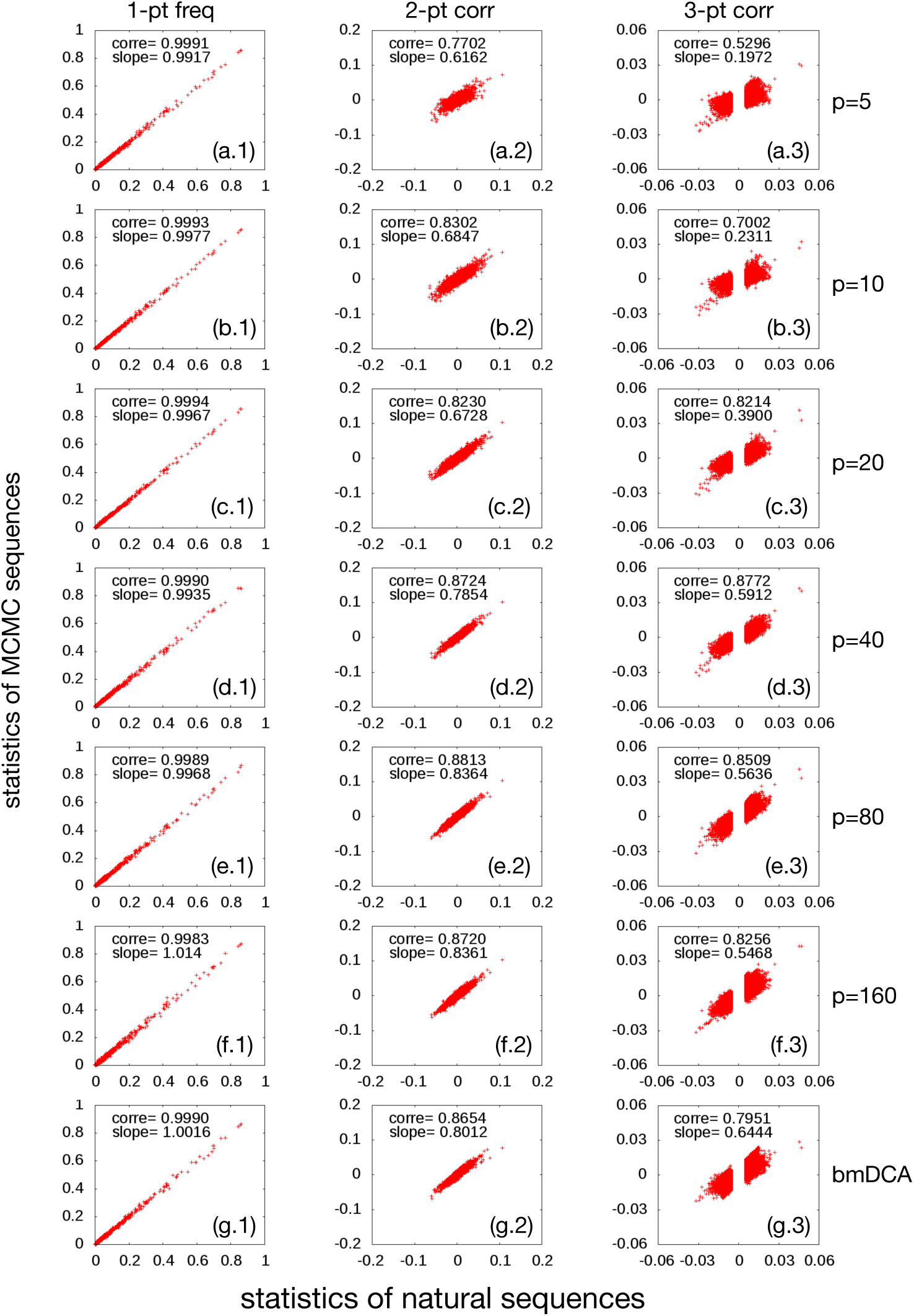
Same as Fig. 2, but for the protein family PF00076.

**FIG. 11:**
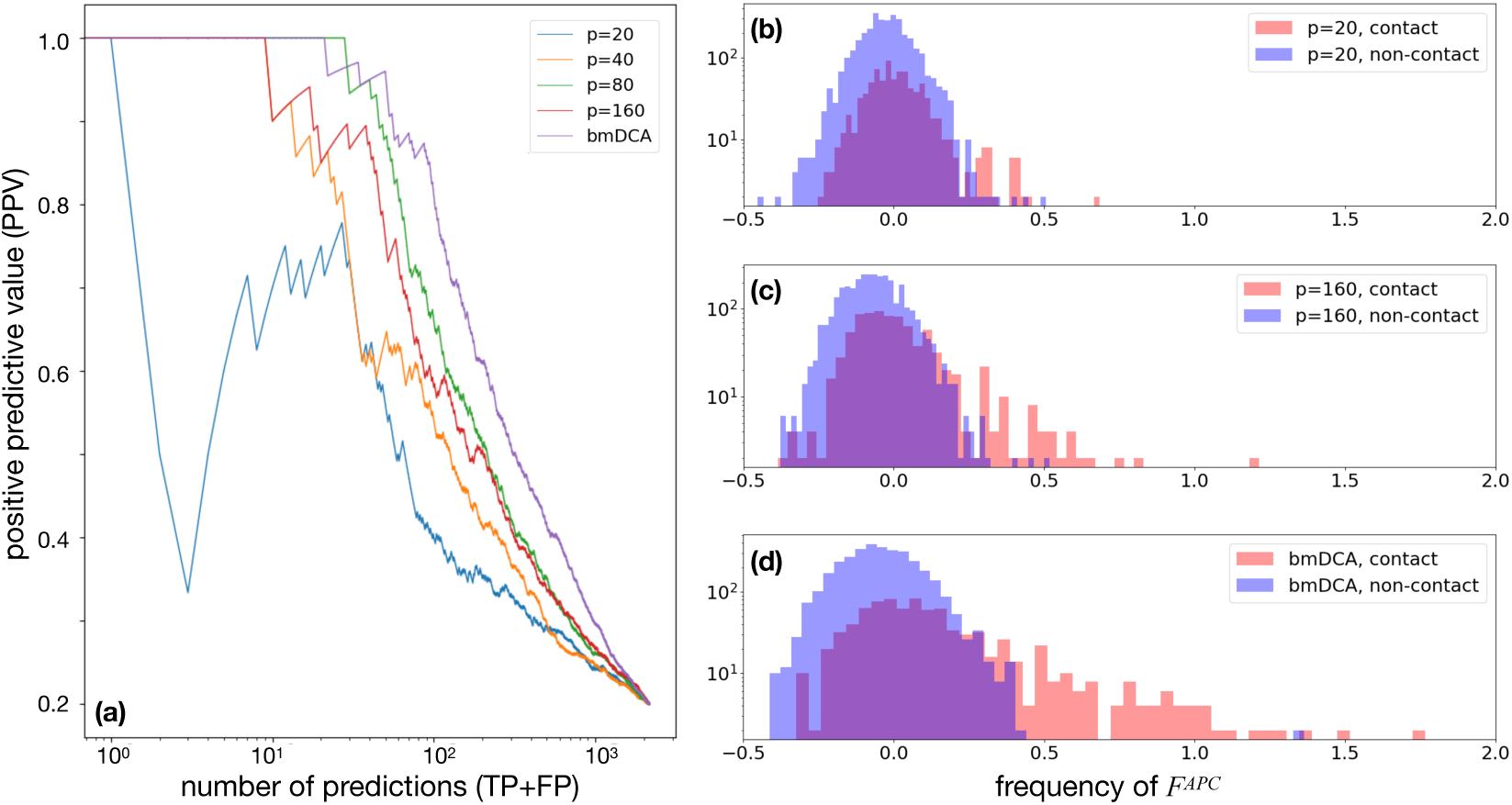
Same as Fig. 3, but for the protein family PF00076.

**FIG. 12:**
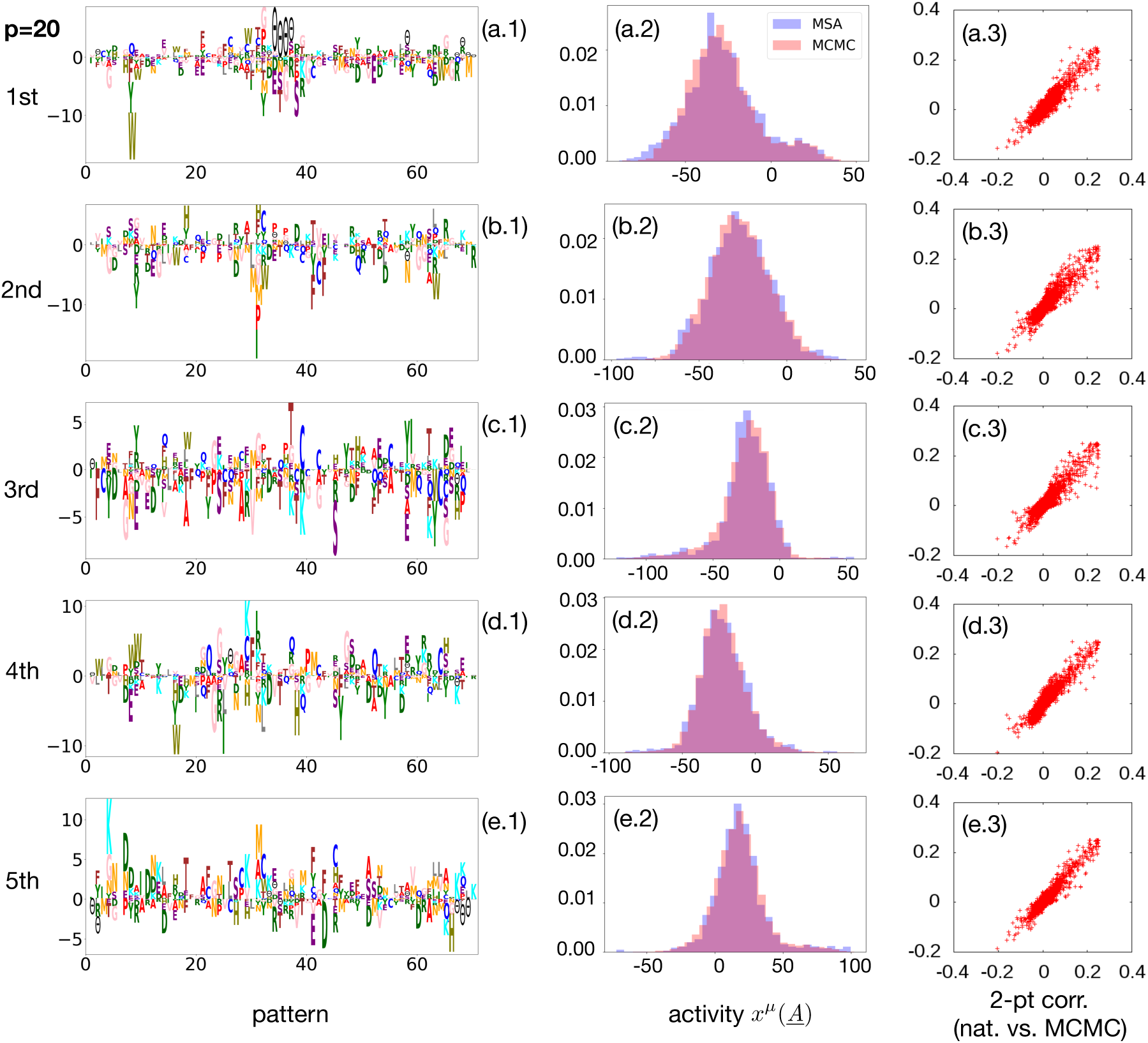
Same as Fig. 5, but for the protein family PF00076.

## Appendix B: Notes and details on inference methods

### 1. Regularization

In case of limited data but many parameters, i.e. the case (Hopfield-)Potts models for protein families are in, the direct likelihood maximisation in Eq. (18) can lead to overfitting effects, causing problems in sampling and parameter interpretation. To give a simple example, a rare and therefore unobserved event would be assigned zero probability, corresponding to (negative) infinite parameter values.

To cope with this problem, regularization is used. Regularization in general penalizes large (resp. non-zero) parameter values, and can be justified in Bayesian inference as a prior distribution acting on the parameter values. In this paper and following [20], we use a block regularization of the form

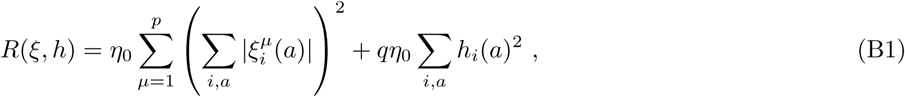

with *η*_0_ being a hyperparameter determining the strength of regularization. This regularization weakly favors sparsity of the patterns.

We use *η*_0_ = *α*_0_*L/qM* with *α*_0_ = 0.0525 as default values throughout this paper. In the last section of this appendix, we show that Hopfield-Potts inference is robust with respect to this choice.

### 2. Contrastive divergence vs. persistent contrastive divergence

#### a. Contrastive divergence does not reproduce the two-point statistics

Contrastive divergence (CD) is a method for training restricted Boltzmann machines similar to persistent contrastive divergence. Initialized in the original data, i.e. the MSA of natural amino-acid sequences, a few sampling steps are performed in analogy to Fig. 1, the *k*th step is used in the parameter update to approach a solution of Eq. (21). However, rather than continuing the MCMC sampling from this sample, the sample is re-initialized in the original data after each epoch. This has, a priori, advantages and disadvantages: The sample remains close to a good sample of the model in CD, but far from a sample of the intermediate model with not yet converged parameters.

As can be seen in Fig. 13, after a sufficient number of epochs the statistics of the CD sample and the training data are perfectly coherent, the model appears to be converged. However, the connected two-point correlations are not well reproduced when resampling the inferred model with standard MCMC. Part of the empirically non-zero correlations are not reproduced and mistakenly assigned very small values in the inferred model.

**FIG. 13:**
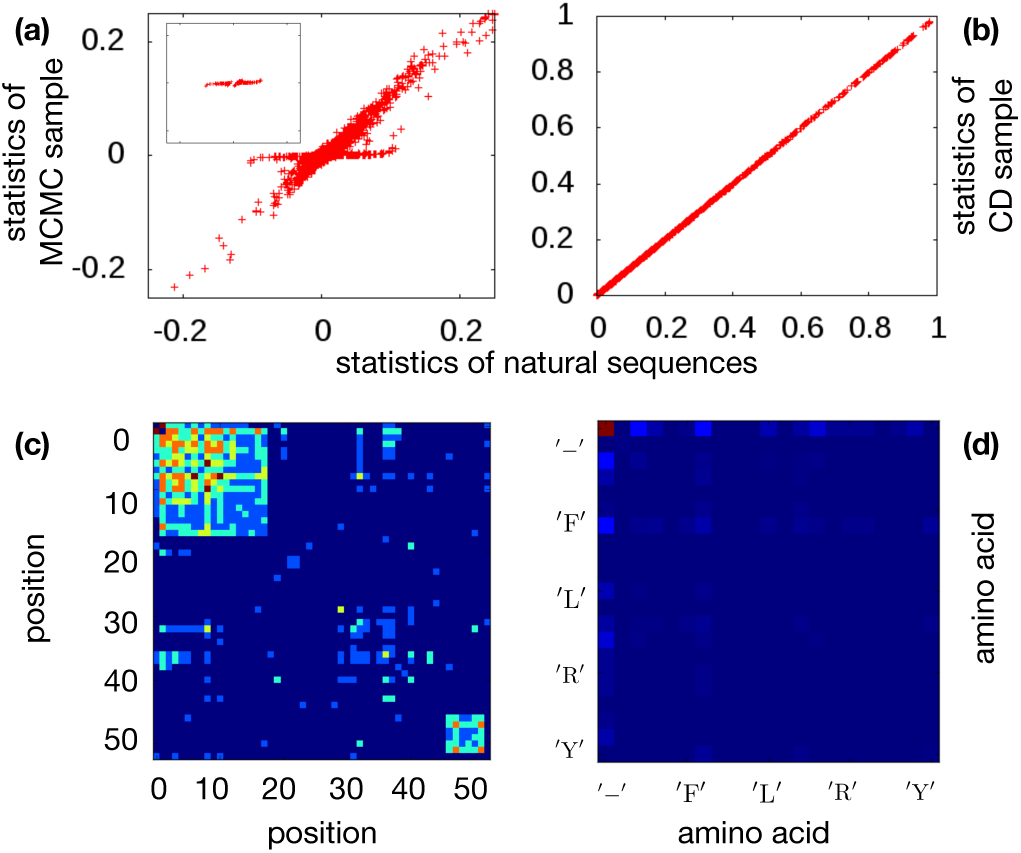
The upper two panels show the statistics (two-point connected correlations) of the training data (PF00014) against an *i.i.d*. MCMC sample extracted from the inferred model, and the CD sample used for inference. The perfect coincidence of the two in the CD case demonstrates that the CD algorithm is converged, contrast is lost despite the fact, that the correlations extracted from an *i.i.d*. sample do not match the empirical ones. To understand this, we have selected those (*i, j, a, b*) with substantial deviations, cf. the insert in the first panel, and analyzed their location in the protein (first panel) and their amino-acid composition (second panel, amino-acids in alphabetical order of one letter code [−, *A, C*, …*Y*]), densities are represented via heatmap plots. Location at the extremities od the sequences and in gap-gap correlations emerge clearly.

To understand this observation, we have selected the elements of the second panel, which show discrepancies between empirical and model statistics, cf. the insert in the figure. The corresponding values of (*i, j, a, b*) are strongly localized in the beginning and the end of the protein chain, and correspond to the gap-gap statistics *c*_*ij*_(−, −). This gives a strong hint towards the origin of the problems in CD-based model inference: gap stretches, which exist in MSA of natural sequences in particular at the beginning and the end of proteins, due to the local nature of the alignment algorithm used in Pfam. Those located at the beginning of the sequence start in position 1, and continue with only gaps until they are terminated by an amino-acid symbol. They never start later than in position 1 or include individual internal amino-acid symbols (analogous for the gap stretches at the end, which go up to the last position *i* = *L*).

In CD only a few sampling steps are performed, so stretched gaps in the initialization tend to be preserved even if the associated gap-gap couplings are very weak. Basically to remove a gap stretch, an internal position can not be switched to an amino-acid, but the gap has to be removed iteratively from one of ts endpoints, namely the one inside the sequence (i.e. not positions 1 or *L*). So, in CD, even small couplings are thus sufficient to reproduce the gap-gap statistics.

If resampling the same model with MCMC, parameters have to be such that gap-stretches emerge spontaneously during sampling. This requires quite large couplings, actually in bmDCA gap-gap couplings between neighboring sites are the largest couplings of the entire Potts model. Using now the small couplings inferred by CD, these gap-stretches do not emerge at sufficient frequency, and correspondingly the positions at the extremities of the sampled sequences appear less correlated.

#### b. Persistent contrastive divergence and transient oscillations in the two-point statistics

Persistent contrastive divergence overcomes this sampling issue by not reinitialising the sample after each epoch, but by continuing the MCMC exploration in the next epoch, with updated parameters.

As shown in the main text, PCD can actually be used to infer parameters, which lead to accurately reproduced two-point correlations when *i.i.d*. samples are generated from the inferred model. However, during inference we have observed transient oscillations, cf. Fig. 14 for an epoch where correlations in a subset of positions and amino acids are overestimated. An analysis of the of the positions and the spins involved in these deviations shows, that again gap stretches are responsible.

**FIG. 14:**
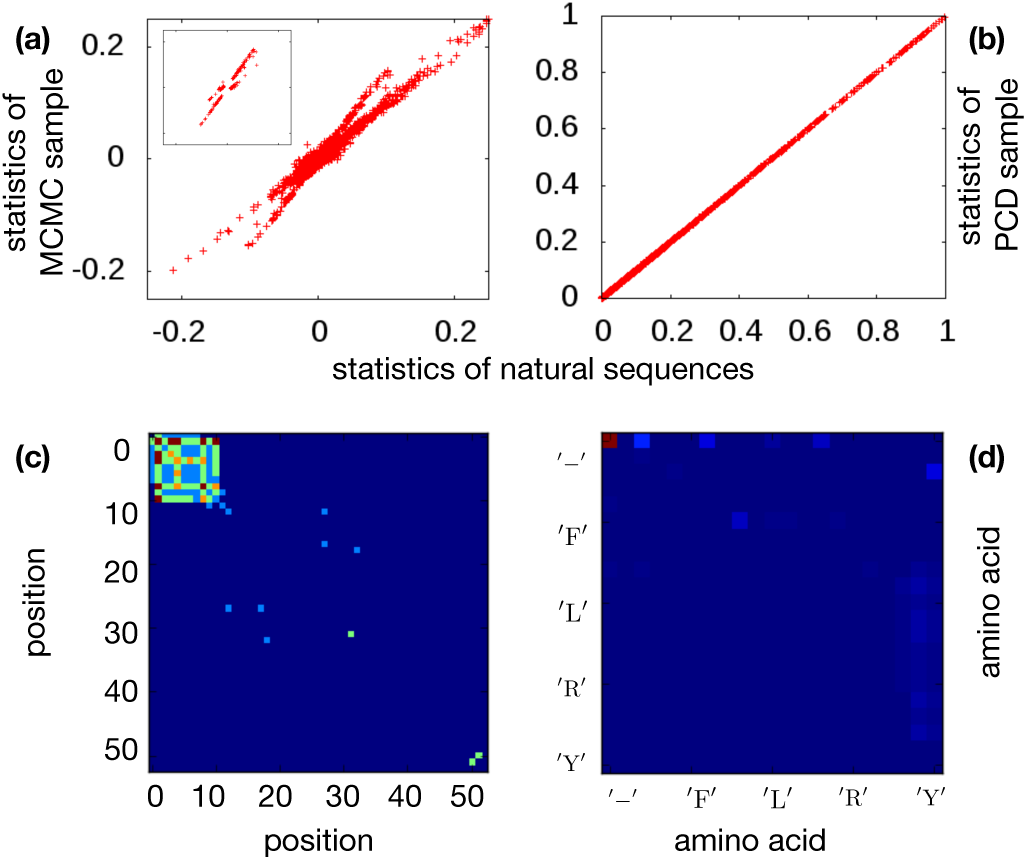
The upper two panels show the statistics (two-point connected correlations) of the training data (PF00014) against an *i.i.d*. MCMC sample extracted from the inferred model, and the PCD sample used for inference. As in the CD case, the MCMC sample shows larger deviations from the empirical observations than the PCD sample, but correlations appear overestimated, and contrast in the PCD plot is sufficient to drive further evolution of parameters. To understand these observation, we have selected those (*i, j, a, b*) with substantial deviations, cf. the insert in the first panel, and analyzed their location in the protein (first panel) and their amino-acid composition (second panel, amino-acids in alphabetical order of one letter code [−, *A, C*, …*Y*]), densities are represented via heatmap plots. Location at the extremities od the sequences and in gap-gap correlations emerge clearly.

The reason can be understood easily. Initially PCD is not very different from CD. Gap stretches are present due to the correlation with the training sample, and only small gap-gap interactions are learned. However, after a some epochs the sample will loose the correlation with the training sample. Due to the currently small gap-gap correlations, gap stretches are lost in the PCD sample. According to our update rules, the corresponding gap-gap couplings will fastly increase. However, due to the few sampling steps performed in each PCD epoch, this growth will go on even when parameters would be large enough to generate gap stretches in an *i.i.d*. sample. Also in the PCD sample, gap stretches will now emerge, but due to the overestimation of parameters, they will be more frequent than in the training sample, i.e. parameters start to decrease again. An oscillation of gap-gap couplings is induced.

The strength of these oscillations can be strongly reduced by removing samples with large gap stretches from the training data, and train only on data with limited gaps. If the initial training set was large enough, the resulting models are even expected to be more precise, since gap stretches do contain no or little information about the amino-acid sequences under study. However, if samples are too small, the suggested pruning procedure may reduce the sample to an insufficient size for accurate inference. Care has thus to be taken when removing sequences.

### 3. Robustness of the results

As discussed before, we need to include regularization to avoid overfitting due to limited data. In Figs. 15 and 16, we show the dependence of the inference results due to changes of the regularization strength over roughly two orders of magnitude. The first of the two figures shows the results for CD: empirical connected two-point correlation are compared with *i.i.d*. samples of the corresponding models. We note that the results depend strongly on the regularization strength. For low regularization, the correspondence between model and MSA is low, due to overfitting. At strong regularization, only part of the correlations is reproduced, we over-regularize and thus underfit the data. For each protein family, and each number *p* of patterns, the regularization strength would have to be tuned.

**FIG. 15:**
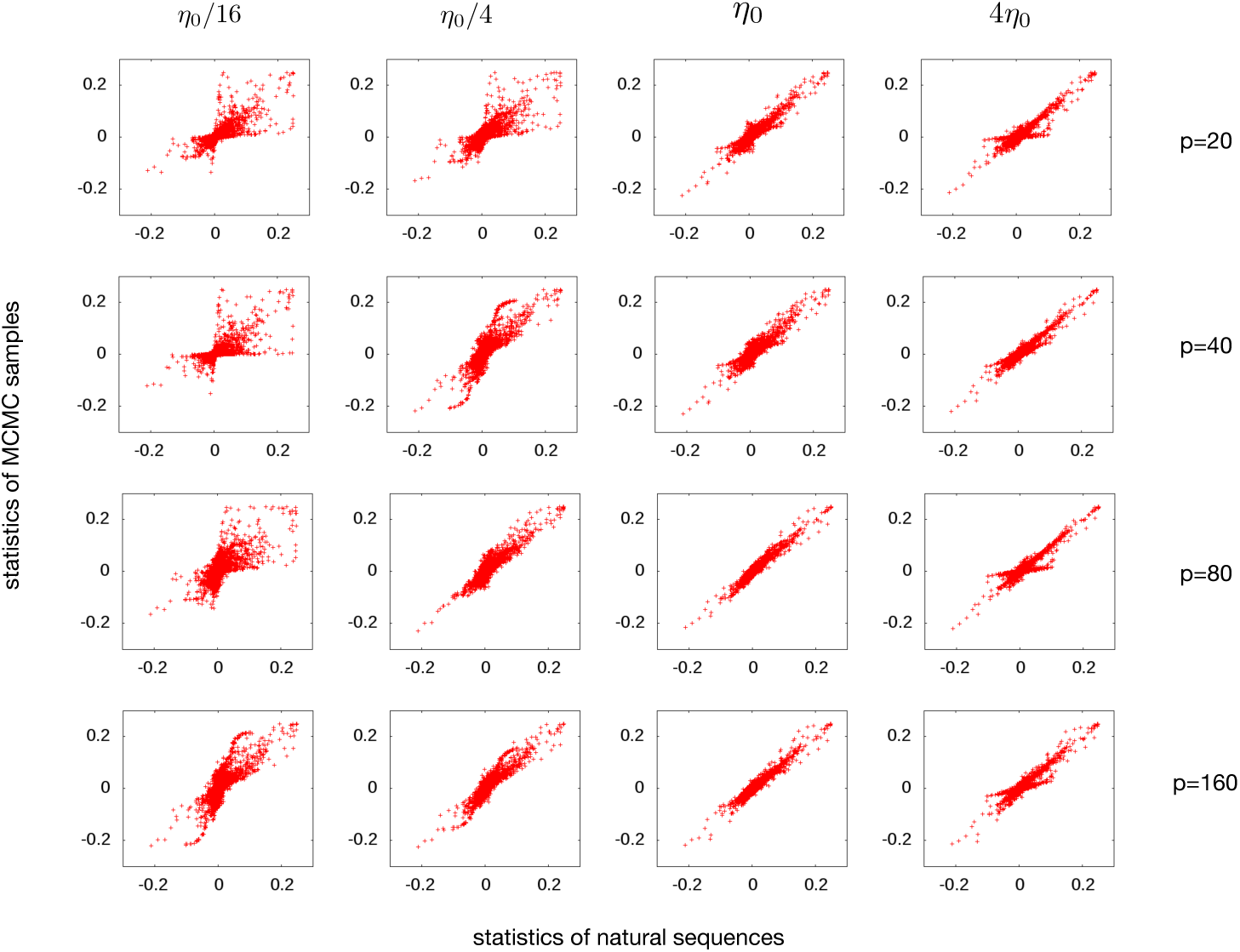
Regularization dependence for CD inference, empirical two-point connected correlations (PF00014) are plotted against those estimated from the model using an *i.i.d*. MCMC sample. The regularization strength is ^0^varied over almost two orders of magnitude, with *η*_0_ = *α*_0_*L/qM* and *α*_0_ = 0.0525, going from a zone of overfitting to one of over-regularization. Results are shown for various values of *p*, illustrating a strong *p*-dependence of the optimal coupling strength.

**FIG. 16:**
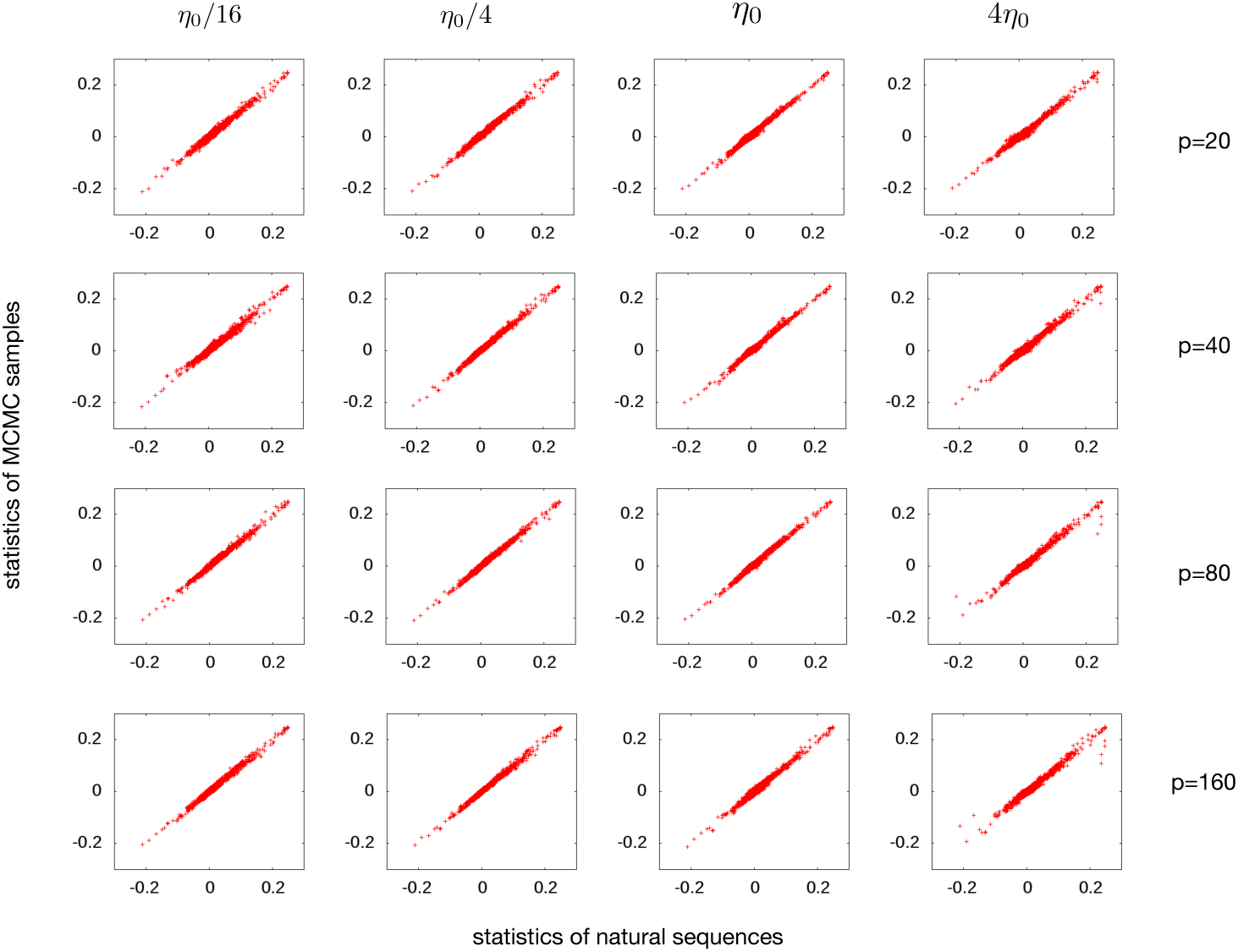
Regularization dependence for PCD inference, empirical two-point connected correlations (PF00014) are plotted against those estimated from the model using an *i.i.d*. MCMC sample. The regularization strength is varied over almost two orders of magnitude, with *η*_0_ = *α*_0_*L/qM* and *α*_0_ = 0.0525; results are shown for various values of *p*. We find a strong robustness of results with respect to regularization.

For PCD, the situation is fortunately much better, results are found to be very robust with respect to regularization, cf. Fig. 16. This allows us to choose one regularization strength across protein families and pattern numbers.

